# Hybrid Genome Sequence of *Cryptococcus neoformans* of Indian origin and Comparative Genome Analysis

**DOI:** 10.1101/2024.08.22.607543

**Authors:** Jananishree Sathiyamoorthy, Jayapradha Ramakrishnan

## Abstract

**Objectives:** The Indian isolate of *Cryptococcus neoformans* underwent complete genome sequencing to elucidate its genomic architecture and functional characteristics. Furthermore, this study aimed to comprehensively characterize the virulence factors (virulome), antibiotic resistance genes (resistome), and the pan-genome of *C. neoformans* spp. through a comparative genomic analysis, providing insights into the genetic diversity and evolutionary relationships among strains.

**Methods:** The genomic data of a clinical strain of *C. neoformans* was assembled and annotated by MaSuRCA5 and Braker tool. Along with this, the assembled genomic data of the 11 strains were retrieved from NCBI datasets. The comparative virulome, resistome, phylogeny and of the 12 *C. neoformans* genomes were analyzed using DFVF, AFRbase, BLAST, CLUSTAL Omega, MEGAX, and Orthovenn3, respectively.

**Results:** The sequenced isolate was identified as a member of the *Cryptococcus neoformans* var. *grubii* subspecies. Notably, virulence-related genes comprise approximately 4.8% of the total genome. A comparative genomic analysis of 12 study genomes revealed variations in virulence patterns, including differences in melanization, immune evasion, blood-brain barrier evasion, transcriptional regulation, and oxidative stress response. The phylogenetic study using MLST and orthologous clusters categorized the subspecies *grubii* and *neoformans* in different clades. Pan-genome analysis showed that 73.6% of orthologous gene clusters and 77.72% of orthologous proteins were conserved across all 12 study genomes, indicating a shared core genome. Furthermore, the evolutionary relatedness study of the pan-genome revealed gene expansion and contraction events among the study strains.

**Conclusion:** This pioneering study presents the first comprehensive genomic and comparative genomic analysis of *Cryptococcus* sp., incorporating data on virulence genes, antibiotic resistance, and pan-genome dynamics. Key findings reveal that strains Cn, H99, and JEC21 harbor crucial virulence genes associated with infection severity. While all study strains possess genes promoting antifungal resistance (AFR), most lack specific single nucleotide polymorphisms defining AFR. Consistent with pan-genome analysis, our results show significant gene expansion and contraction events in these strains. This study underscores the importance of bioinformatic tools for efficient whole-genome analysis and large-scale comparative genomics research.

## 1. Introduction

*Cryptococcus* sp. is a dimorphic environmental fungus that majorly infects HIV patients [1]. It is considered intricate because it is classified into two broad species, *C. neoformans* and *C. gatti*. Furthermore, two subspecies, *C. neoformans* var*. grubii* (serotype A) and *C. neoformans var. neoformans* (serotype D) were found. *C. gatti* was classified into serotype B and serotype C. Currently, there are nine major molecular types [2]. Among these, serotype A (*C. neoformans var. grubii*) is highly pathogenic and the molecular types VNI and VNII are widespread in the Indian population [3]. *C. neoformans* is considered one of the critical priority pathogens by WHO [4]. The high pathogenicity of serotype A is due to its escalated virulence factors, environmental adaptability, and immune evasion capabilities [5,6].

This ubiquitous pathogen frequently inhabits avian environments (pigeon droppings, claws and wings of pigeons), marshy areas, and decaying wood [7,8]. The infection begins in the lungs through the inhalation of cryptococcal spores/ infectious propagules. Once *Cryptococcus* colonizes the lungs it breaches the lungs and circulates to the liver, kidney and bone marrow through the bloodstream and disseminates to the brain via paracytosis, transcytosis and trojan horse mechanism [9,10]. The *Cryptococcus* sp. exhibits neurotropism owing to the presence of proteinases and urease that allow them to penetrate the blood-brain barrier (degradation of ECM components of the host) and survive. Tropical, subtropical and temperate locations have been found to have higher incidences of cryptococcal infection [11]. The prevalence of cryptococcal infection was high majorly in Sub-Saharan Africa (incidence: 6-10% among individuals with advanced HIV disease; mortality: 24% to 47% at 10 weeks post-diagnosis), followed by Asia and Pacific (incidence: 1 to 3% among the HIV infected individuals) [12]. These areas are influenced by a combination of environmental, ecological, socioeconomic, and healthcare-related factors. It was estimated that 1, 52, 000 new cases and 1, 12, 000 (19%) deaths were reported globally each year. The advent of HAART reduced the incidence of cryptococcal infection but the mortality rate remains high [12].

In the recent decade, it was noted that immune-competent individuals are also acquiring the infection and the reason could be the result of the organism’s increased pathogenicity, which could be caused by virulence, stress tolerance, or Anti-Fungal Resistance (AFR) [9]. The virulence factors of *Cryptococcus* sp. are the capsular polysaccharides (glucoronoxylomannan, galactoxylomannan), melanin, mannoproteins and certain lytic enzymes (urease, phospholipase and proteases) [13,14].

The preliminary role of the stress tolerance genes was to overcome the environmental stressors (temperature, pH, reactive oxygen species, reactive nitrogen species and nutrient starvation). In addition, the autophagy genes (also stress-tolerant genes) in *Cryptococcus* sp., play a role in cell differentiation, cell growth, morphology and pathogenesis. These ATGs play a critical role in bilateral reproduction [15–17]. The recent insights of cryptococcal genomics reveal that stress-tolerant genes play a pivotal role in modulating the major virulence factors synthesis and promoting AFR [18]. They also regulate the genes responsible for maintaining the cell wall, cell membrane integrity and also regulate the evasion of host immune response.

The development of AFR or tolerance, which is brought about by stable and heritable point mutations that impact the target and drug interaction is significantly influenced by the organism’s genetic background [19]. Complete Genome Sequencing (CGS) has revolutionized the understanding of AFR in *C. neoformans.* Though the inherent resistance (echinocandins resistance) is more common in *Cryptococcus* sp. the fungal genomics highlights the insights of acquired resistant mechanisms (target protein modification, over-expression of target proteins, and upregulation of multi-drug transporters) caused by genetic mutations and structural variations[19–21]. The mutation’s non-primary target proteins also produce resistance to antifungal drugs [22].

In the earlier decades, PCR-Restriction Fragment Length Polymorphism and PCR fingerprinting were used to study *Cryptococcus* molecular epidemiology, the genetic structure of the fungus and genome plasticity. The recent advent and advancements in WGS help in understanding the epidemiology and genetic structure of *Cryptococcus* sp.[23]. Globally the studies based on the WGS have demonstrated small genetic variability and a small number of chromosomal rearrangements across the molecular types and also the variations in gene content and structural changes due to translocations, aneuploidy and inversions were noted in *C. neoformans*. The WGS also helps in revealing the hybrids present between the *Cryptococcus* sp., complex [23]. In addition, WGS helps in identifying the microevolution signals of infecting strains revealing the rapid adaptation of the organism through hypermutation to identify genes related to the adaptation of an organism to the stress condition that include virulence and pathogenicity features and monitoring of outbreaks [23,24].

Considering it as a critical priority pathogen as described in the guidelines by WHO and the gap in the knowledge about the genomic features of clinical *Cryptococcus* species of Indian origin, this was the first study in India we demonstrated hybrid sequencing of *Cryptococcus neoformans* (clinical strain) and unravel the genomic aspects and comparative pan-genome analyses were made with the clinical isolates and an environmental isolate from NCBI datasets to understand the evolutionary relationships, resistome and virulome. To the best of our knowledge, this is the first kind of study to perform the hybrid sequence of the *Cryptococcus* genome, its resistome, virulome and pan-genome analysis.

## 2. Materials and Methods

### 2.1. Culture Isolation and DNA extraction

The comprehensive methodology is depicted in Figure 1. The *Cryptococcus* sp. (Cn) was collected from the microbiology lab of Trichy Medical College, Tamil Nadu, India sourced from the cryptococcal meningitis patient. The presumptive identification was performed by Indian ink staining after growing the yeast colonies in potato dextrose agar containing streptomycin. The purified yeast colonies were stored in 30% glycerol at –80℃.

**Figure 1:**
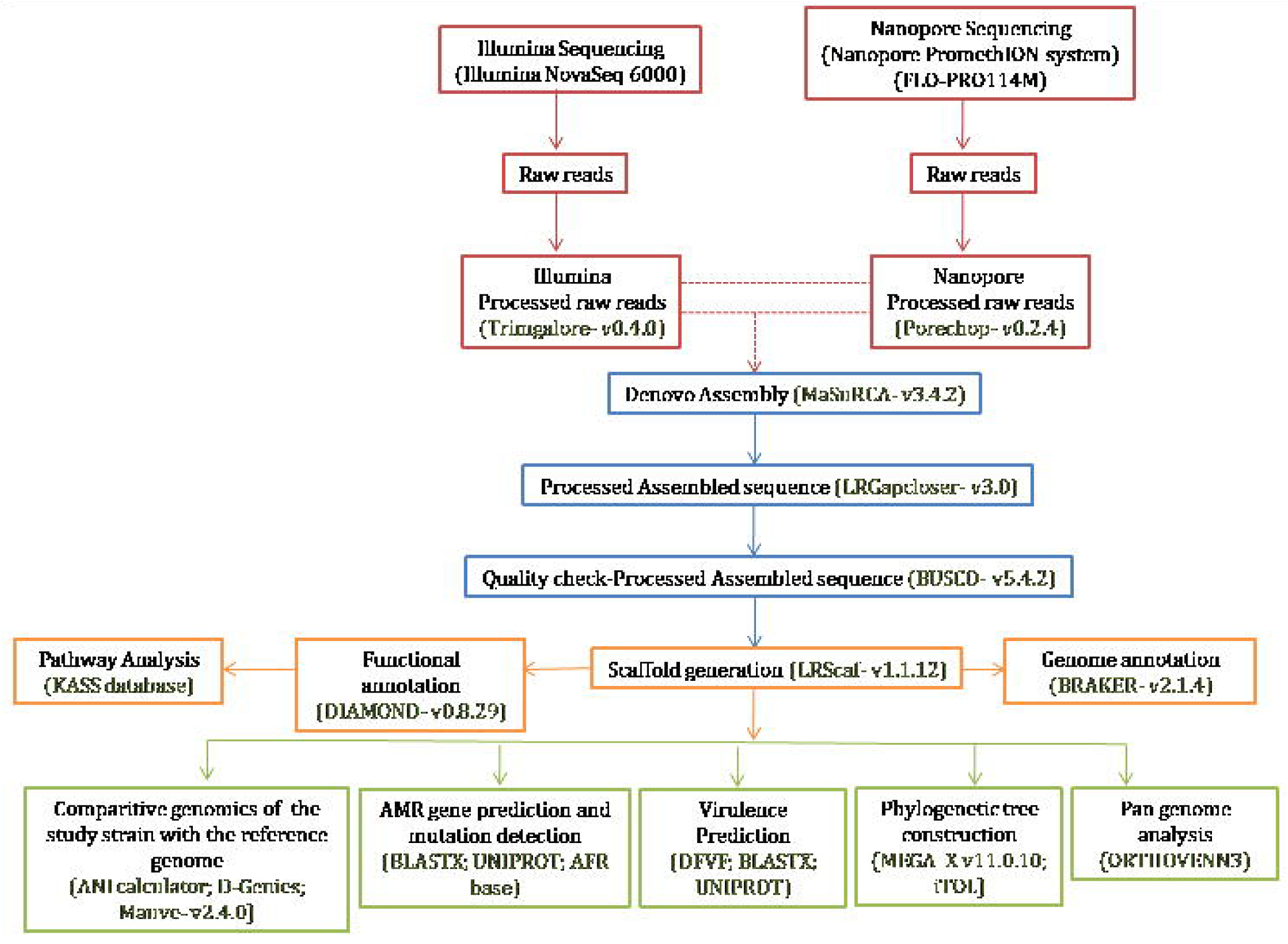
A flow chart on the comprehensive methodology used in the study.

Initially, the clinical isolate Cn was grown in Potato Dextrose Broth (PDB) and incubated for 48 hours at 37°C. Approximately 10^7^ log phase cells were obtained by centrifugation at 7500 rpm for 10 minutes following the scheduled incubation period. The isopropanol precipitation technique, phenol-chloroform, and urea lysis buffer were used to extract the DNA from the pellet. After two cycles of mechanical lysis, one cycle was finished with one milliliter of urea lysis buffer and four hours of incubation at 30°C. Proteinase K (0.02 ml) and Sarkosyl lysis buffer (0.4 ml) were added to the lysate, and the mixture was incubated for 1 hour at 55°C. After a three-minute break on ice and another three minutes of mechanical lysis at 2500 rpm, the cycle was repeated three times. RNase A was added to the lysate, and it was then incubated for 20 minutes at 65°C. The lysate was blended with an equal volume of phenol-chloroform, inverted, and centrifuged for 10 minutes at 4°C and 13,200 rpm. After transferring the aqueous phase to a fresh microcentrifuge tube and adding an equivalent volume of chloroform, isoamyl alcohol was added and the tube was centrifuged. After transferring the aqueous phase to a fresh microcentrifuge tube, DNA was precipitated in an equivalent volume of isopropanol. The pellet was washed with 70% ethanol, air dried and dissolved in 10mM Tris-Cl (pH 8.0).

The concentration and purity of genomic DNA were quantified using a Nanodrop Spectrophotometer (Thermo Scientific; 2000). DNA integrity was observed on agarose gel electrophoresis (supplementary figure 1). DNA concentration quantified using Qubit dsDNA HS assay kit.

### 2.2. Complete Genome Sequencing

#### 2.2.1 Illumina library preparation, sequencing and preprocessing

The library preparation was performed with 50ng of quantified DNA. The QIASeq FX DNA Library Kit was used to prepare the library according to the protocol provided in the kit manual[25]. Indexing PCR was performed on the library to enrich the adapter-tagged fragments and purification of the amplified library was performed with High Prep TM beads. The library was quantified using a Qubit fluorometer (Thermo Fisher Scientific, MA, USA), and the Agilent 2200 Tape Station was used to evaluate the fragment size distribution of the library (supplementary figure 2).

The libraries were sequenced on Illumina NovaSeq 6000 (Illumina, San Diego, USA) using 150 bp paired-end chemistry. The data obtained from the sequencing run was de-multiplexed using Bcl2fastq2 softwarev2.20 and FastQ files were generated based on the distinct dual barcode sequences. The sequencing quality was assessed using FastQC3 v0.11.8 software and Trimgalore-v0.4.0.

#### 2.2.2. Nanopore Library preparation, Sequencing and preprocessing of reads

The NEBnext ultra II end repair kit was used to end polish and A-tail the genomic DNA. A blunt TA ligase master mix was used to barcode the end-prepared samples. With NEB Quick T4 DNA Ligase, sequencing adapters were ligated onto double standard DNA fragments to create the library. The prepared library was ready to use the following purification using AMPure XP beads. Qubit was used to quantify the prepared library.

The estimated DNA was combined and sequenced using a PromethION flow cell (FLO-PRO114M) on a Nanopore PromethION system (PromethION P24 and Data Acquisition Unit, ONT, Oxford, UK) following the necessary data amount and effective library concentration. Porechop-v0.2.4 was used to process the Nanopore raw reads.

#### 2.2.3. Assembly and Annotation

The high-quality processed reads were used for d*e novo* assembly using the MaSuRCA5-v3.4.2 assembler [26]. The assembled genome was further improved by closing the gaps using LRGapcloser6-v3.0 and finally, scaffolds were constructed using the LRScaf-v1.1.12 [27]. The assembled genome was utilized for gene and protein prediction using the eukaryotic genome prediction tool, Braker-v2.1.4 gene prediction program [28]. The predicted proteins were searched against the Uniprot fungal database using DIAMOND-v0.8.29 [29]. The KAAS database was used to retrieve information from the BlastP program for functional annotation and pathways [30]. The database uses BLAST comparisons against the manually edited KEGG GENES database to offer functional annotation of genes [30]. Along with the annotation of the isolated study strain Cn, 11 more assembled sequences were also annotated by using the tool AUGUSTUS v3.4.0 [31].

### 2.3. Comparative genome analysis with reference genome

The assembled genome was also compared with the reference genome using two web-based resources, the Average Nucleotide Identity (ANI) calculator and D-Genies which are tools used to quickly compare the sequences at the whole genome scale [32]. The assembled genome was validated for completeness and quality of the genome assembly using BUSCO-v5.4.2 [33]. The Mauve-v2.4.0 tool was used to compare the draft genome against the closest reference genome for the construction of multiple genomes in the presence of large-scale evolutionary events such as rearrangement and inversion [34].

### 2.4. Retrieval of global genome sequence data of *Cryptococcus neoformans*

The WGS of 11 isolates of *C. neoformans* were collected from the NCBI datasets (<u>https//</u>www.ncbi.nlm.nih.gov/datasets/) in FASTA format. The sequences were further analyzed for an evolutionary relationship, virulence, AFR and pan-genome. Due to the lack of fungal databases for virulence and AFR analysis, we have limited our comparative study to 12 isolates (n=11 isolates from NCBI datasets and n=1 hybrid sequenced genome).

### 2.5. Phylogenetic analysis

The phylogenetic analysis was carried out between the study strain Cn and the retrieved *C. neoformans* from the NCBI datasets. The extracted sequences from NCBI datasets include the species isolated from clinical (n=10) and environment (n=1). To investigate the phylogenetic placement based on MLST, the seven housekeeping genes *CAP59*, *GPD1*, *IGS1*, *SOD1*, *LAC1*, *URA5* and *PLB1* were used. Each genome was concatenated and the phylogenetic tree was constructed by the maximum likelihood method using MegaX software. Tree editing and annotation were performed using the interactive Tree of Life (iTOL) [35].

### 2.6. Virulence gene identification and comparative genomic study

The virulence-related genes (VRGs) (n=272) of *C. neoformans* were collected from the literature and DFVF (Database of Fungal Virulence Factors) database [15,16,18,36–43]. The listed gene sequences were retrieved from Uniprot and their presence in the annotated study genomes were identified using BLAST. The sequence with 99 to 100% identity was considered for the presence or absence of VRGs. The comprehensive result of the comparative virulome analysis was consolidated using iTOL.

### 2.7. AFR gene identification and comparative Analysis of Mutation Profiles

Similarly, the AFR genes of *C. neoformans* were collected from the literature. The AFR gene sequences were retrieved from Uniprot and their presence in the annotated genome was identified using BLAST with the query coverage and identity of 99 to 100% is considered as the presence or absence of virulence genes. Followed by, the protein mutations responsible for the antifungal resistance provided in the AFRbase database were compared with the annotated genomes of the study strains using CLUSTAL Omega. The comprehensive result of the comparative AFR analysis and mutation profiles was consolidated using iTOL.

### 2.8. Pan-genome analysis

The pan-genome analysis was performed using Orthovenn3 with a FASTA annotation file and GFF file produced by AUGUSTUS (genome annotation tool). The data was analyzed and visualized in Orthovenn3. The orthologous gene analysis was carried out by the OrthoFinder algorithm and the Markov clustering algorithm was used to generate the gene clusters. The evolutionary phylogenetic tree construction was made using FastTree2 by the maximum likelihood method. The gene family contraction and expansion analysis were performed using CAFE5.

## 3. Result

### 3.1. Sequencing and Assembly

The hybrid technology resulted in a total of 7.35 million reads at 116.05X coverage for Illumina sequencing and a total of 6.09 lakhs raw reads at 104.8X coverage for Nanopore sequencing. The genome sequences of the strain Cn were deposited under BioProject accession number PRJNA650119. The assembled genome was approximately 19 MB in size with 20 contigs. The quality check of the assembled genome showed 89.6% completeness.

### 3.2. Gene prediction and annotation

A total of 5673 predicted proteins were annotated successfully using the BRAKER tool. The gene ontology annotation categorizes the proteins into 3 groups such as biological process (14.87%), cellular components (41.75%) and molecular function (43.37%). The detailed gene ontology annotation is provided in Figure 2a. The functional annotation using KEGG pathways annotated 5963 genes which fall majorly into 6 categories, which include BRITE hierarchy (3186 genes), metabolism (1294 genes), genetic information processing (831 genes), cellular processes (419 genes) and environmental information processing (97 genes). The genes (n=136) of unknown functions are categorized separately. The assembled genome was identified with 29 cellular pathways using the KAAS database. The top 10 functional pathways are depicted in Figure 2b.

**Figure 2:**
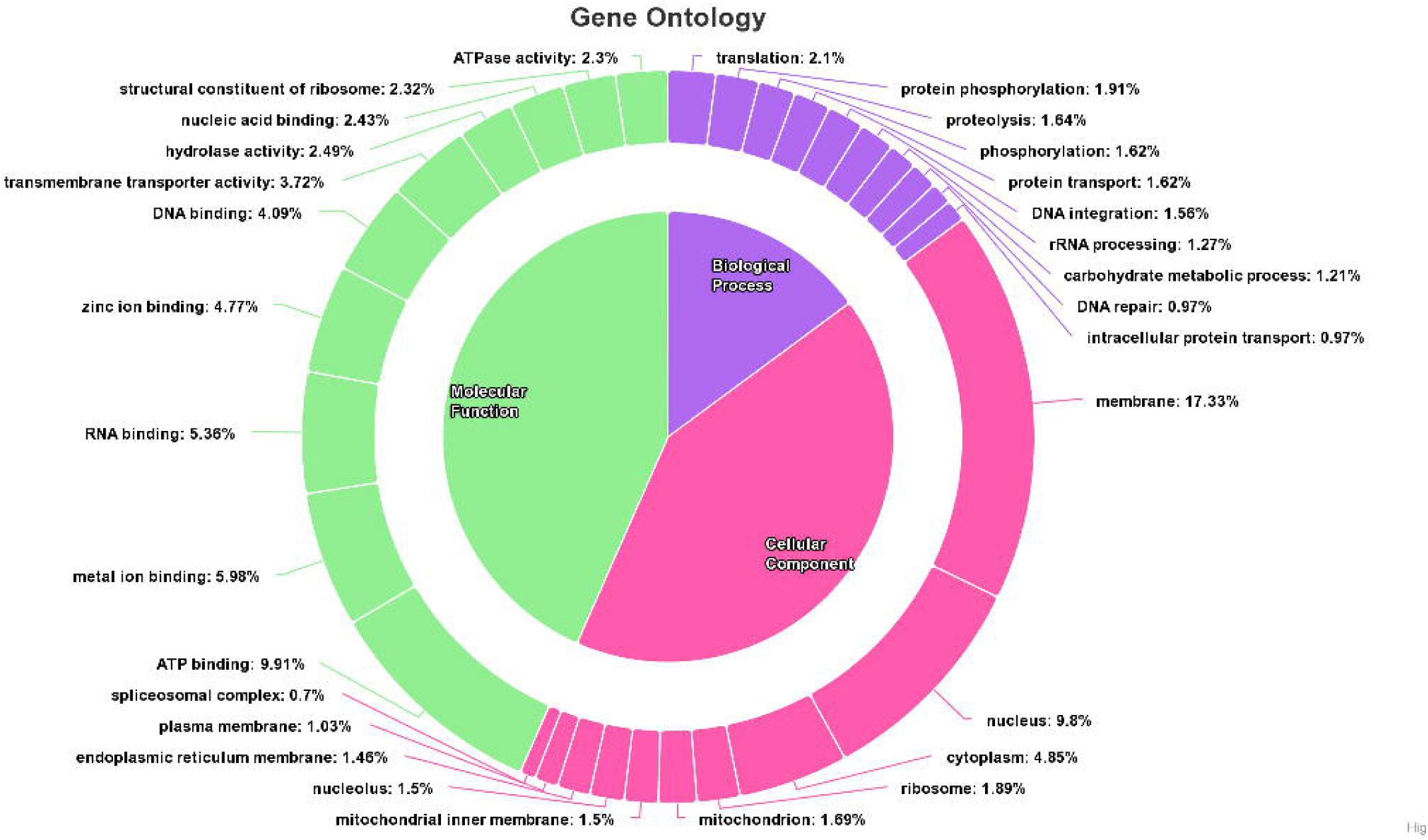
Functional annotation: (a) GO annotation by DIAMOND database: the Gene ontology of the study strain Cn and (b) Pathway analysis by KAAS database: Top 10 pathway functions of the study strain Cn.

### 3.3. Genome identity search of clinical Cn

The genome identity search using the ANI calculator indicated a 99.9% identity with the reference genome (H99). The same result was depicted in a whole genome analysis by MAUVE as a dot plot image and D Genies by a linear pairwise genome comparison image (figure 3a, 3b and 3c). The comparison of ITS region was shown to have 100% coverage and identity with *C. neoformans* var. *grubii* CBS8710 strain, isolated from Hodgkin’s lymphoma patient from North Carolina.

**Figure 3:**
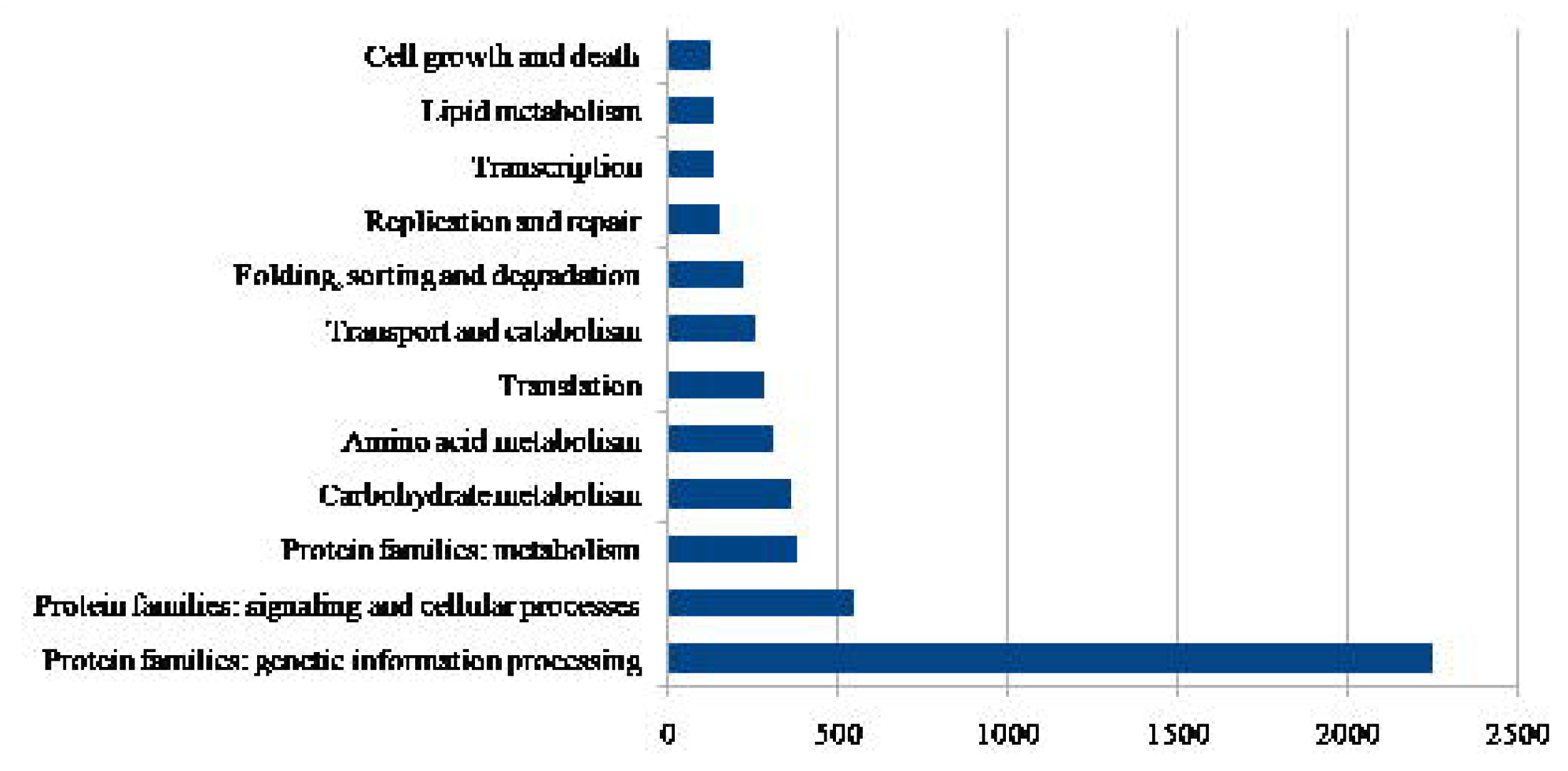
Genome identity search: (a) ANI Calculation of the study strain Cn and reference strain H99; (b) Dot plot comparison using D GENIES and (c) Linear genome image using MAUVE.

### 3.4. Phylogenetic analysis using MLST

The 12 study genomes were analyzed for phylogeny by MLST genes using the MEGAX and visualized using iTOL. Phylogenetic analysis showed that all the South African isolates were grouped under the same single clade, whereas isolates from USA, India and Australia were grouped into another clade. The phylogenetic tree showed that the clinical and environmental isolates were indistinguishable, and the subspecies *grubii* and *neoformans* were distinguishable. The midpoint of the tree was divided into two major clades. **Clade 1**: Clinical strain (JEC21) from the USA, belonging to subspecies *neoformans.* **Clade 2**: Clinical and environmental strains belonging to subspecies *grubii*, further divided into: **Subclade 1**: Environmental strain (VNII) from the USA. **Subclade 2**: Clinical strains (Bt65, Bt81, Bt89, Bt133) from South Africa, grouped into 2 clusters. **Subclade 3**: Clinical strains from India (Cn), Australia (KBCN0140, KBCN0142, SACN00B1), USA (C23, H99), grouped into 5 clusters (Figure 4)

**Figure 4:**
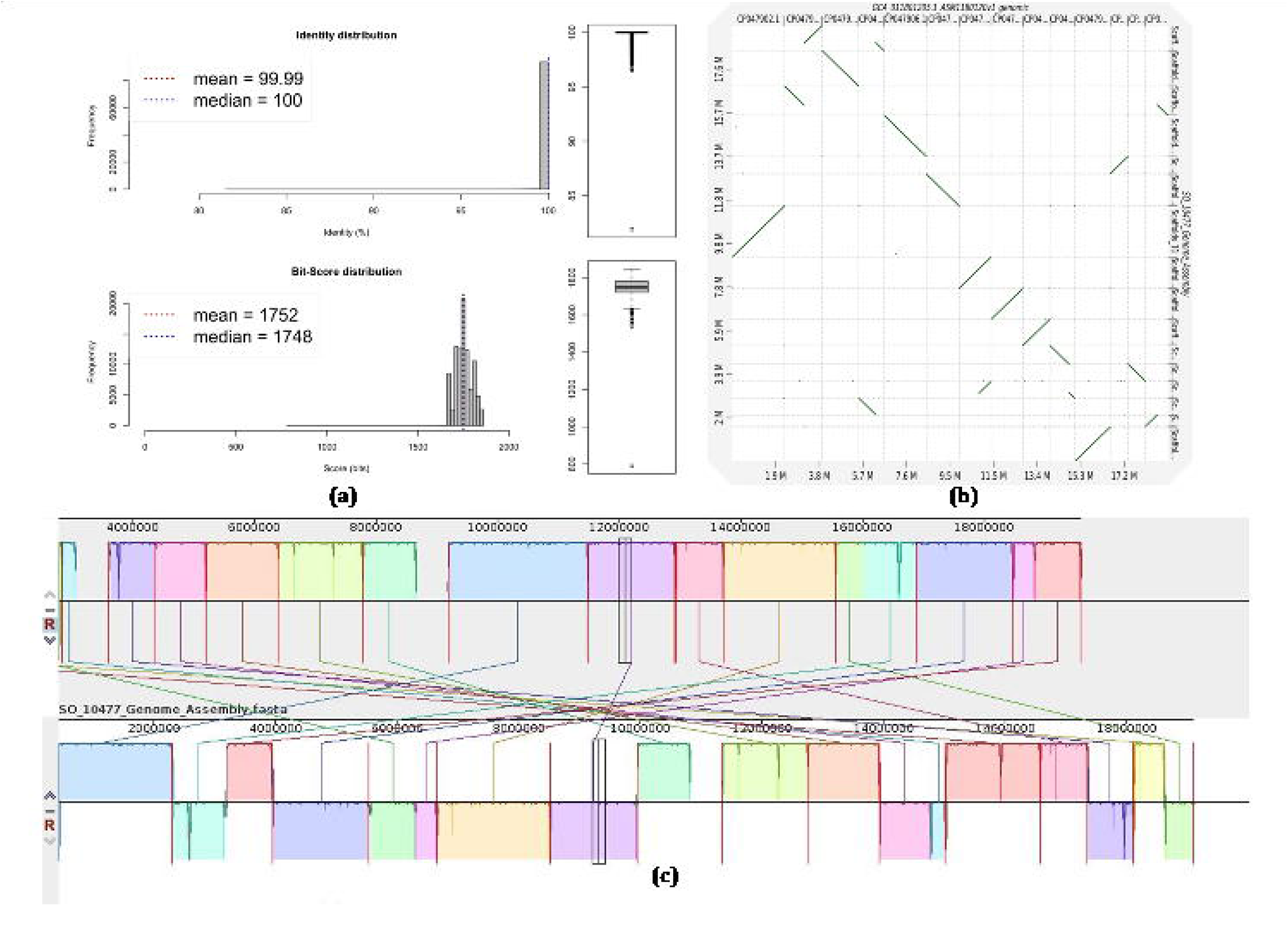
Phylogenetic tree based on MLST genes: MLST phylogeny was constructed using the maximum likelihood method with MegaX. 1000 bootstrap replicates were calculated to assess robustness. The midpoint of the phylogenetic tree was divided into two major clades by the subspecies *grubii* (red line) and *neoformans* (yellow line). The tree picture represents the environmental strain and the human represents the clinical strain. The magenta colour represents the Botswana (South Africa) strains, the blue colour represents the TamilNadu (India) strain, the orange colour represents the Melbourne (Australia) strains, the violet colour represents the Tennessee (USA) strain and the red colour represents the California (USA) strains.

### 3.5. Virulome and comparative virulome analysis

Due to the lack of virulome profiling databases, the Virulence Related Genes (VRGs) collected from the literature and DFVF database were identified in the 12 genomes using BLAST (>99% coverage and >99% identity). From the literature and DFVF database, 272 VRGs of *C. neoformans* were retrieved. Based on the functionality we categorized the VRGs into stress tolerance (n=164), capsular-associated (n=78), cell wall-associated (n=85), cell membrane-associated (n=24), nutrient acquisition, iron acquisition and homeostasis (n=39), host cell adhesion (n=32), host cell colonization (n=16), evasion of host immune response (n=49), biofilm-associated (n=9), protease (n=7), urease (n=1), toxin production (n=14), melanin associated (n=33), titan cell-associated (n=6) and morphogenesis related (n=13) genes. Thus, most of the genes involved in virulome analysis are multifaceted either directly or indirectly involved in more than one virulence trait. The list of genes in the study strain Cn belonging to each category is briefed in Figure 5.

**Figure 5:**
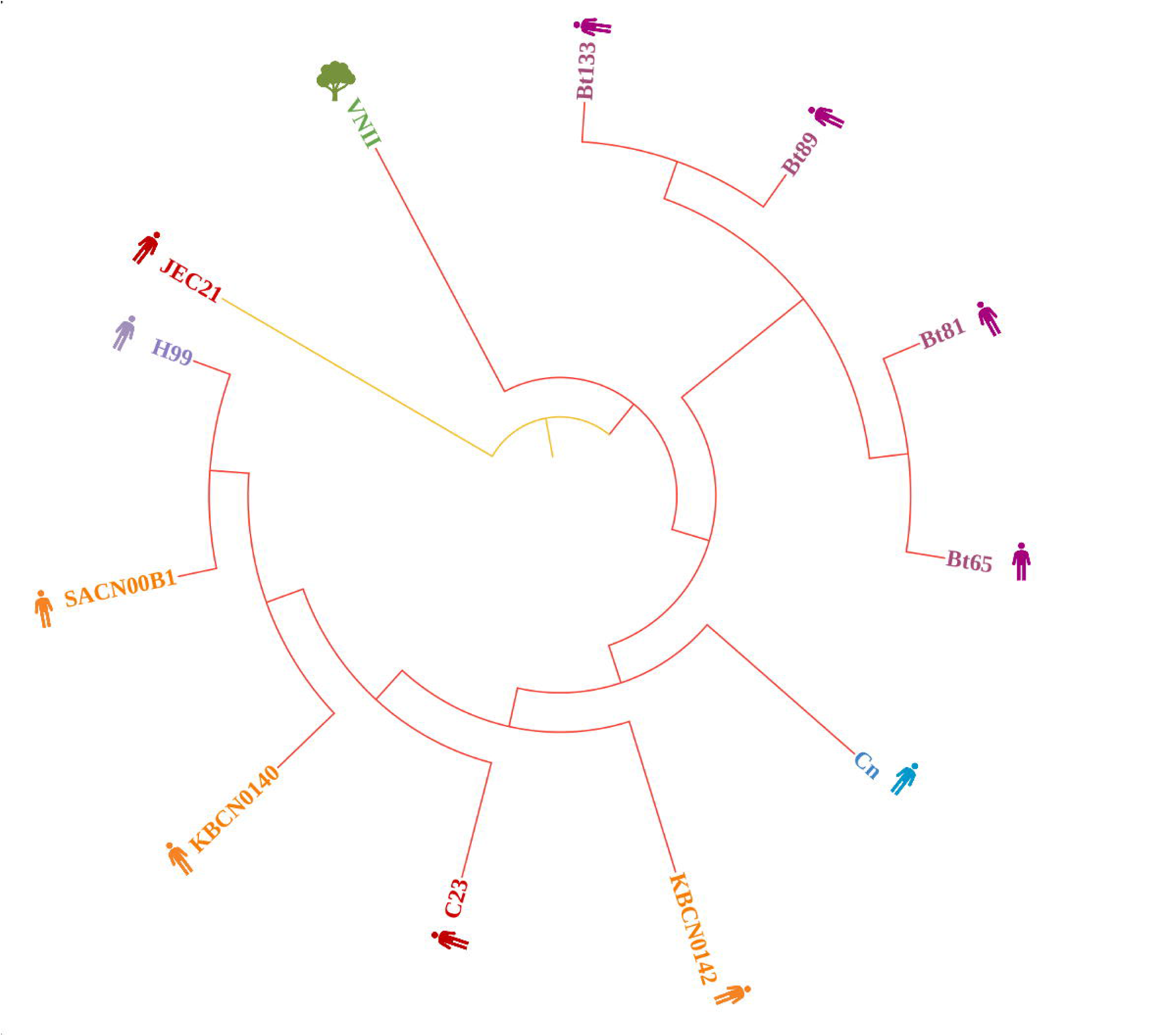
VRGs present in the study strain Cn: The chart represents the (n=264) VRGs of Cn that were categorized into stress tolerance, cell wall-associated, cell membrane-associated, capsule-associated, morphogenesis associated, cell adhesion, immune evasion, iron acquisition, protease, nutrient acquisition, copper homeostasis, host cell membrane disruption and melanin associated genes.

In the virulome comparative study, H99 has the highest % of virulence genes with 99.26% (n=270) and the lowest in Bt133 with 82.14% (n=224). However, both are clinical isolates. The Cn strain has 97.06% (n=264) of virulence genes. Being an environmental isolate, VNII has 83.45% (n=227) virulence genes. For the comparative virulome analysis, the major VRGs polysaccharide capsule (*CAP10*, *CAP59*, *CAP60* and *CAP64*), melanin (*LAC1* and *LAC2*), thermotolerance (Hsp90 and *STU1*), cell adhesion (*ALG3*), iron acquisition (*CFT1*, *SIT1*, *CIR1* and RIM101), oxidative stress tolerance (*SOD1* and *SOD2*) and enzymes (*PLB1*, *URE1*, *YME1*, *RIM13*, *APR1*, *PIM1*, *SPP1*, *RBD1* and *APP2*) were used. The comparison revealed that these genes were found in almost all the *C. neoformans* spp. It was found that the strains Cn, H99 and JEC21 have all 24 major VRGs and could be more virulent than other strains (figure 6; table 1). The clinical strains Cn and H99-infected animal models have also proven to be highly virulent.

**Figure 6:**
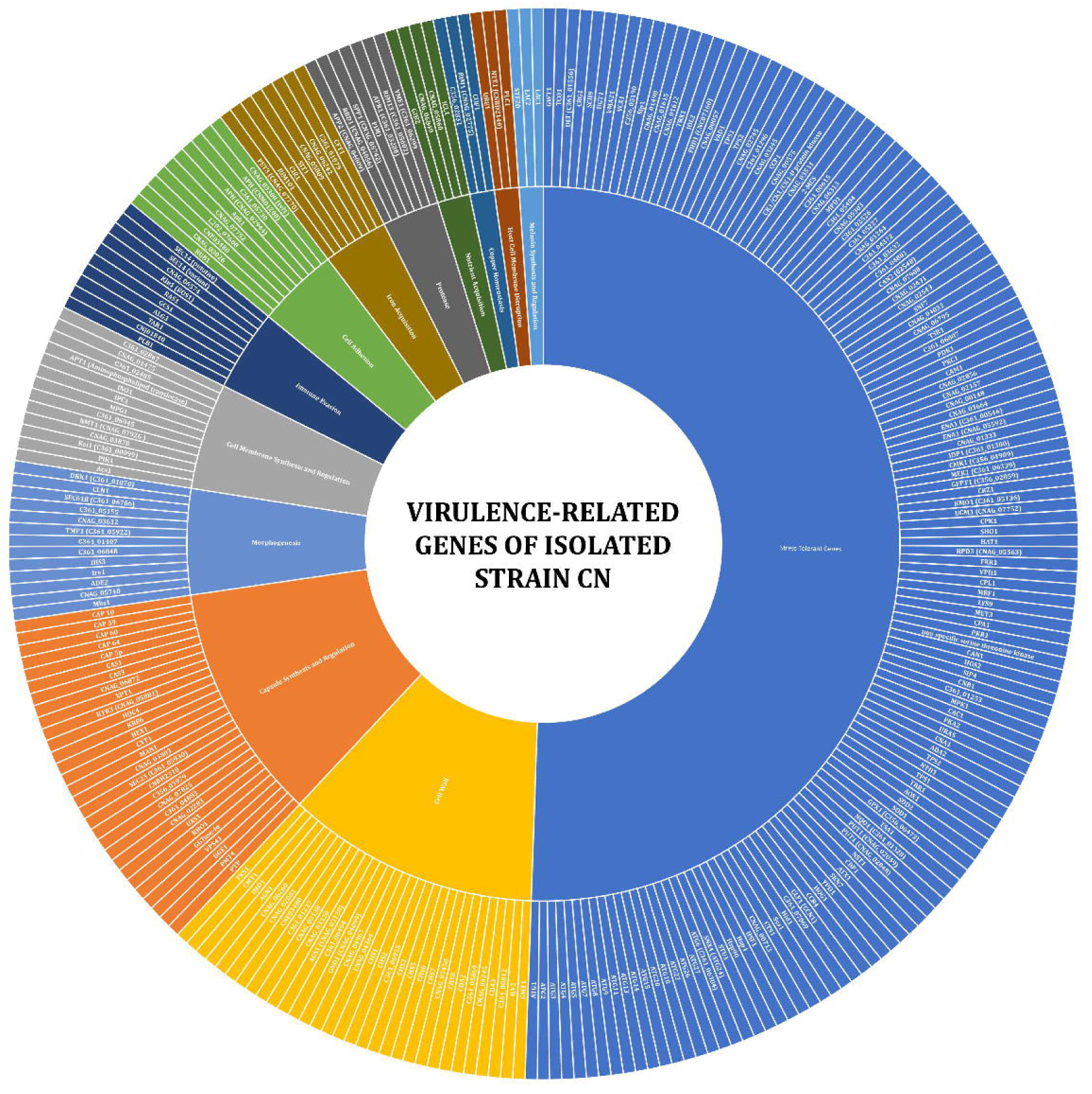
Comparative virulome analysis: The graph represents the comparative study of the major VRGs present in the (n=12) study strains. The colored region represents the presence of genes.

**Table 1:**
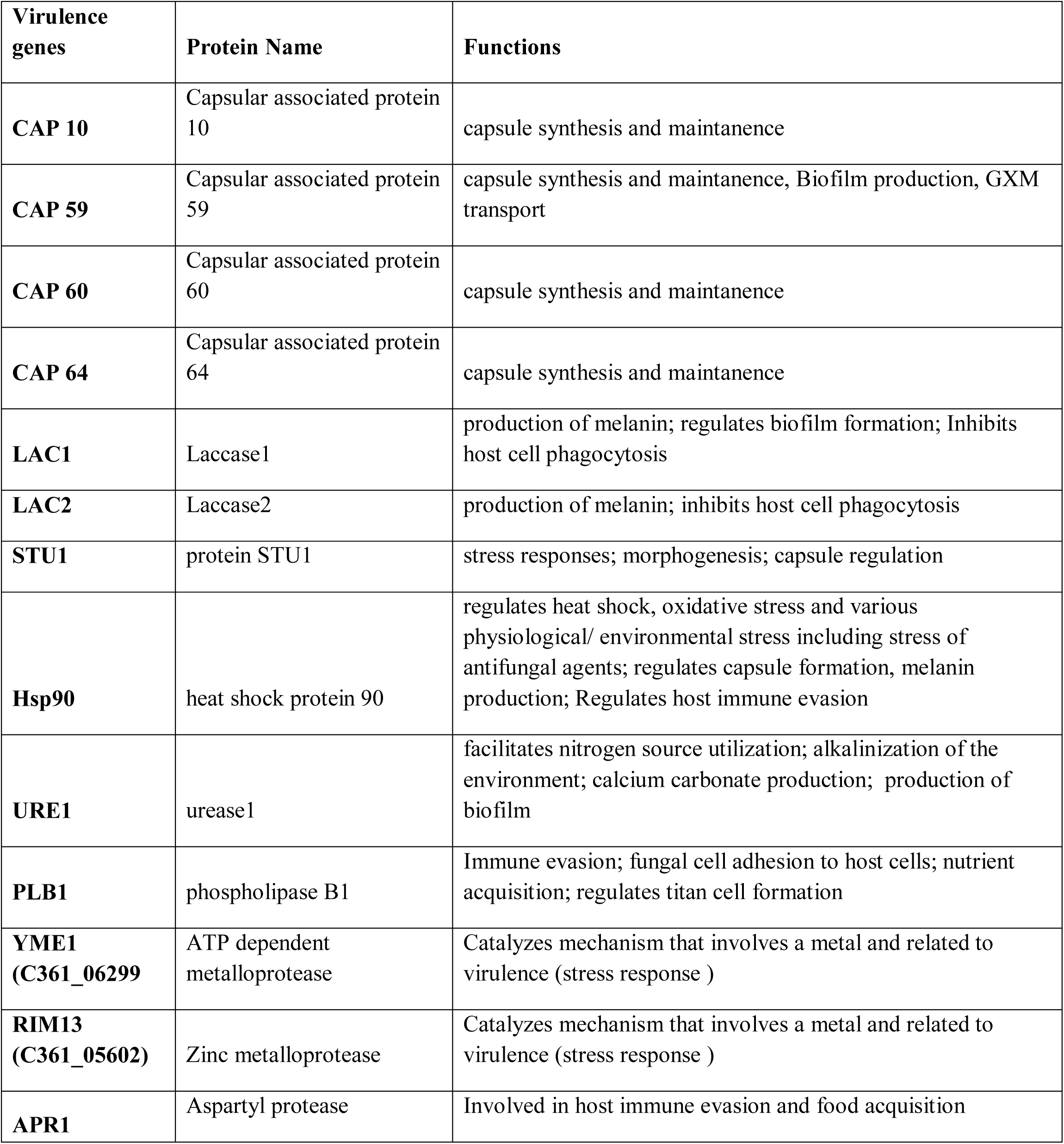
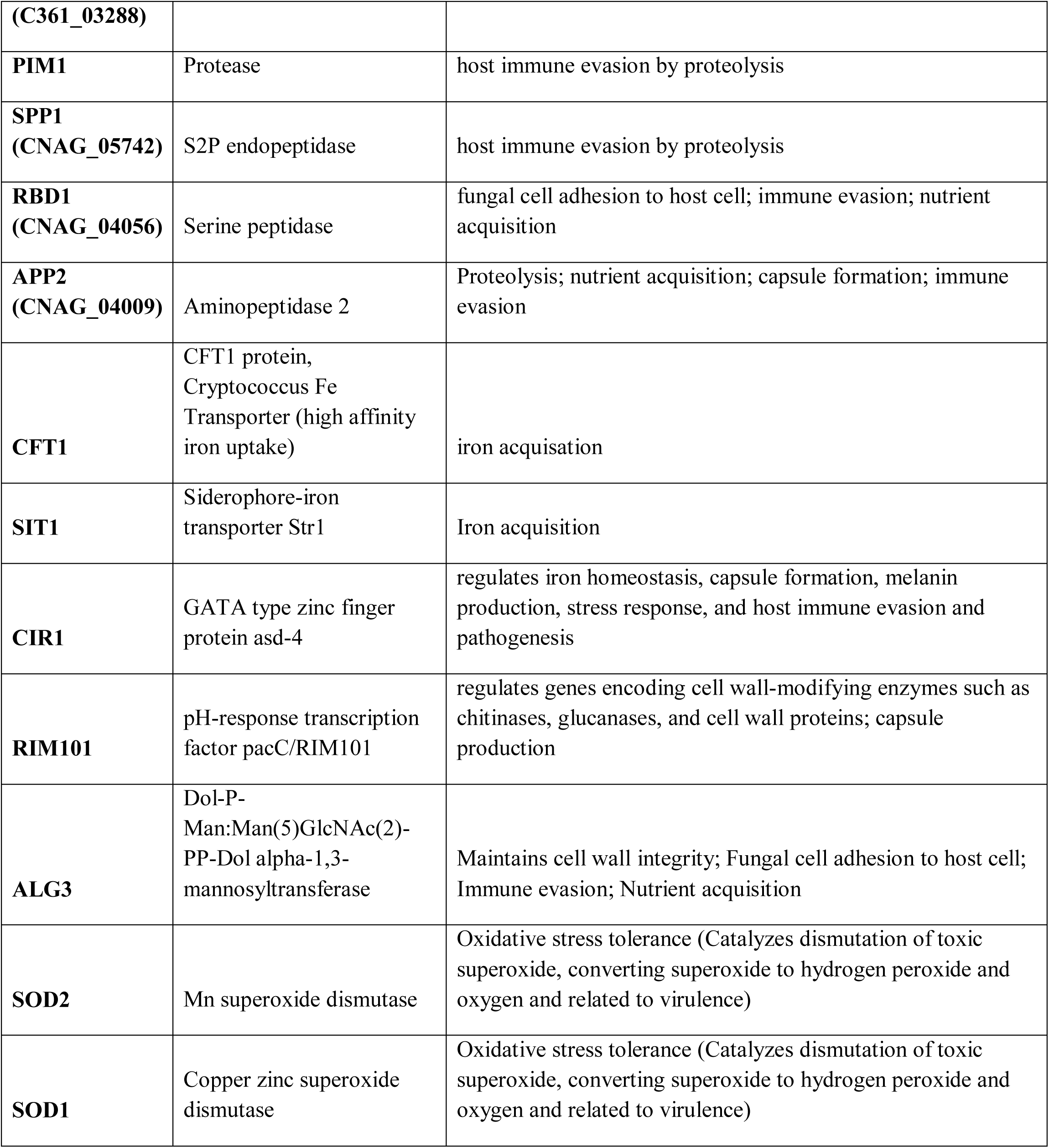
The major VRGs of *C. neoformans* study strains.

### 3.6. Resistome and Comparative Resistome Analysis

The AFR genes retrieved from the literature are examined for their presence in the study genomes using BLAST. The output significantly predicted AFR genes with >99% identity and sequence coverage. Around 78 genes were predicted and observed as factors promoting antifungal resistance. The Cn has ABC transporters (n=6 genes), Major Facilitator Superfamily (n=13), drug target genes (n=12), Inherent resistance genes (n=2), Stress regulator genes (n=2), DNA mismatch genes (n=5) and transcriptional regulator genes (n=37) contributing to AFR (figure 7).

**Figure 7:**
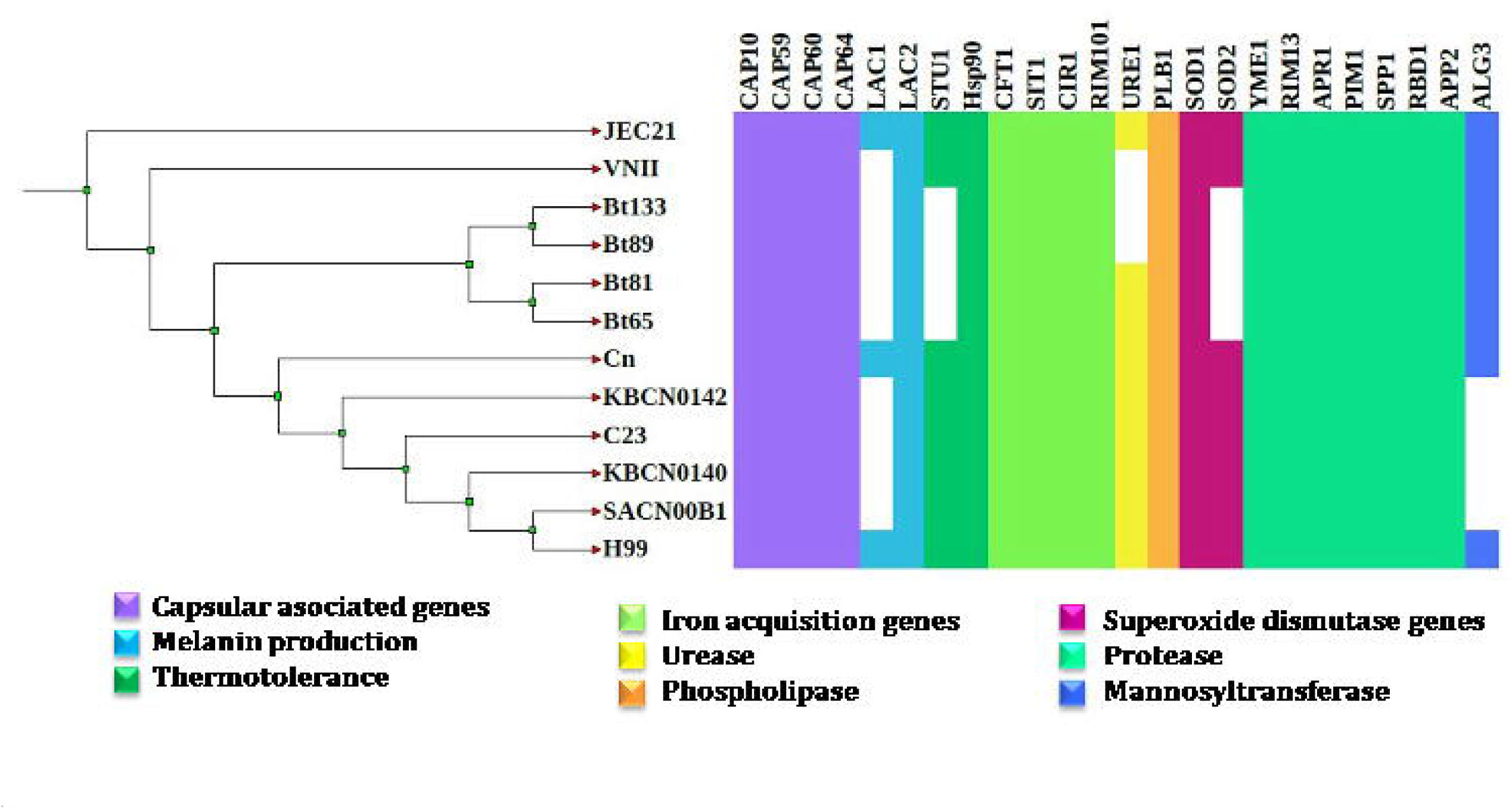
AFR genes present in the study strain Cn: The chart represents the (n=78) AFRs of the Cn that were categorized into transporter, structural, DNA mismatch, nucleic acid synthesis, stress response and transcription factors.

Among the 12 strains, the Cn was found to harbor abundant AFR genes across certain classes of antifungals. Resistance to azole drugs were contributed by *ERG11, ERG24, ERG1, SRE1, AFR1, AFR2, PMR5, TUB1, SEY1, YOP1, TUB1* and *YAP1* genes. Resistance to echinocandins was contributed by *FKS1, CRM1,* and *Hsp90* genes. The gene for P4 ATPase and *CRM1* confers intrinsic resistance for echinocandins and the resistance to pyrimidine analog was conferred by the genes *FCY2*, *TMP1* and *FUR1.* Allylamines resistance was provided by the gene *ERG1.* Conveying that the strain Cn (n=78; 100%) has more AFR genes than other isolates. The possible reason would be that the hybrid genome has been identified to have more AFR genes than other strains. Despite being an environmental strain, the VNII has 53 AFR genes (68.83%).

Among 20 major antifungal resistance genes, 13 genes were commonly shared (100%) in all the 12 study strains. The genes *ERG11*, *ERG24*, *SEY1* and *ERG1* belonging to sterol biosynthesis provide resistance by the alteration of target protein structure, overexpression of target protein and increased activation of efflux pump. The gene for P4-ATPase and the gene *CRM1* provide inherent resistance by regulating the intracellular calcium concentration. The gene *FKS1* (β-glucan synthase; Cell wall synthesis) and Hsp90 (Heat shock protein 90; Cell wall regulation) provide resistance by alteration, under expression of the target protein and stress regulation. The genes *TMP1* and *FUR1* (Thymidylate synthase 1 and Uracil phosphoribosyltransferase 1; nucleotide biosynthesis) provide resistance by alteration and overexpression of target proteins. The gene *PMR5* (ABC transporter PMR5; ABC transporter) is the only common transporter protein that provides resistance to antifungal drugs (figure 8).

**Figure 8:**
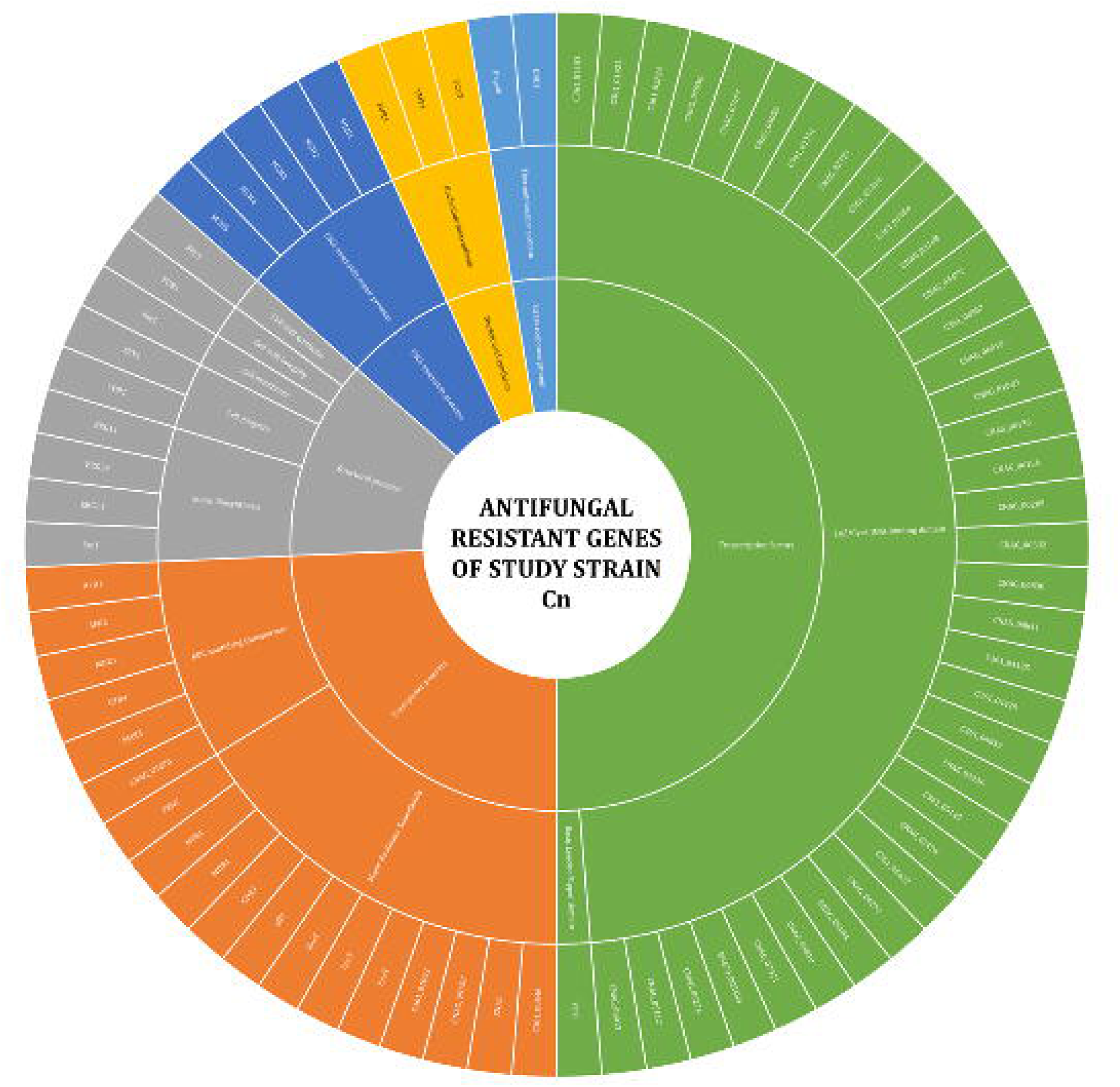
Comparative resistome analysis: The graph represents the comparative study of the major AFR genes present in the (n=12) study strains. The colored region represents the presence of genes.

Among the 78 AFR genes, SNPs were detected in *ERG11* (lanosterol 14-α demethylase), *FKS1* (β-glucan synthase), *UXS1* (UDP-glucuronate decarboxylase 1), *FUR1* (Uracil phosphoribosyltransferase), *TUB1* (β tubulin) and *FCY2* (Cytosine permease) genes. Among them, *ERG11* was noted to have the highest number of mutations (n=9), followed by *UXS1* (n=2), *FKS1* (n=1), *FUR1* (n=1) and *FCY2* (n=1). The presence of SNPs in the 12 study strains is depicted in Figure 9; Table 2. In the *FKS1* gene, serine at the 643^rd^ position was mutated to proline. This resulted in the conformational change in the lipid enriched molecules, as its role is to anchor the lipid molecules, and amino acid serine at 643^rd^ position is involved in the binding site. In the *ERG11* gene phenylalanine (F) at 126^th^ position has been changed to serine (S) and glycine at 464^th^ position has been mutated to valine. Both these residues were not conferred to active site binding pocket. In the *TUB1*, histidine at the 6^th^ position was changed to glutamine, glutamic acid at 98^th^ position was mutated to glycine and phenylalanine at 200^th^ position was mutated to asparagine. Glycine at the 190^th^ position was mutated to glutamic acid in the ABC transporter gene *FUR1*. In all the 12 study genomes, though the mutation was observed, none of them were conferred to the active site binding pocket which is also concurrent with our *in vitro* susceptibility test where the study clinical strain Cn showed sensitivity to fluconazole and amphotericin B [44].

**Figure 9:**
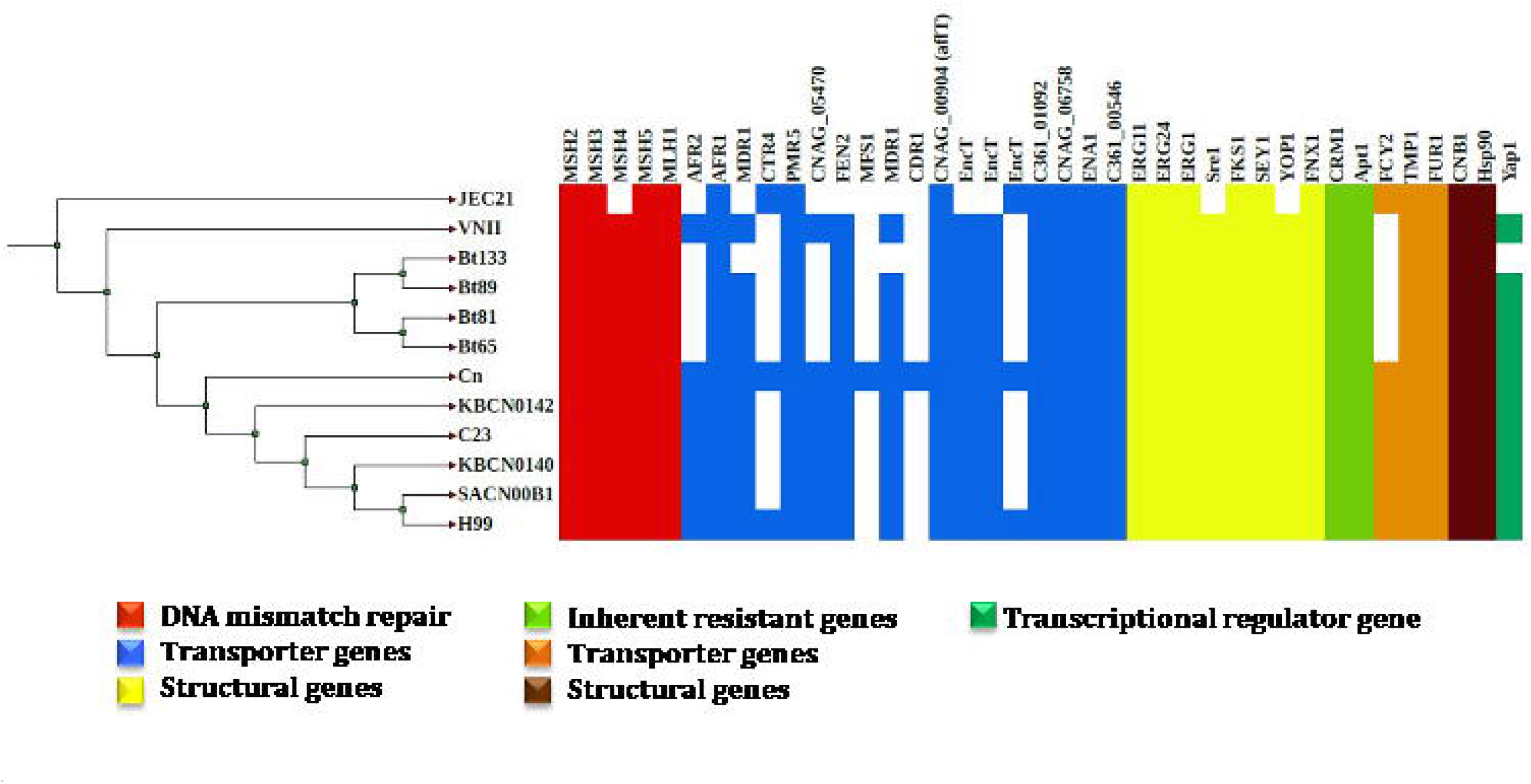
Comparative analysis of SNPs in the AFR genes: The graph represents the comparative study of the SNPs present in the AFR genes of the (n=12) study strains. The colored region represents the presence of genes.

**Figure 10:**
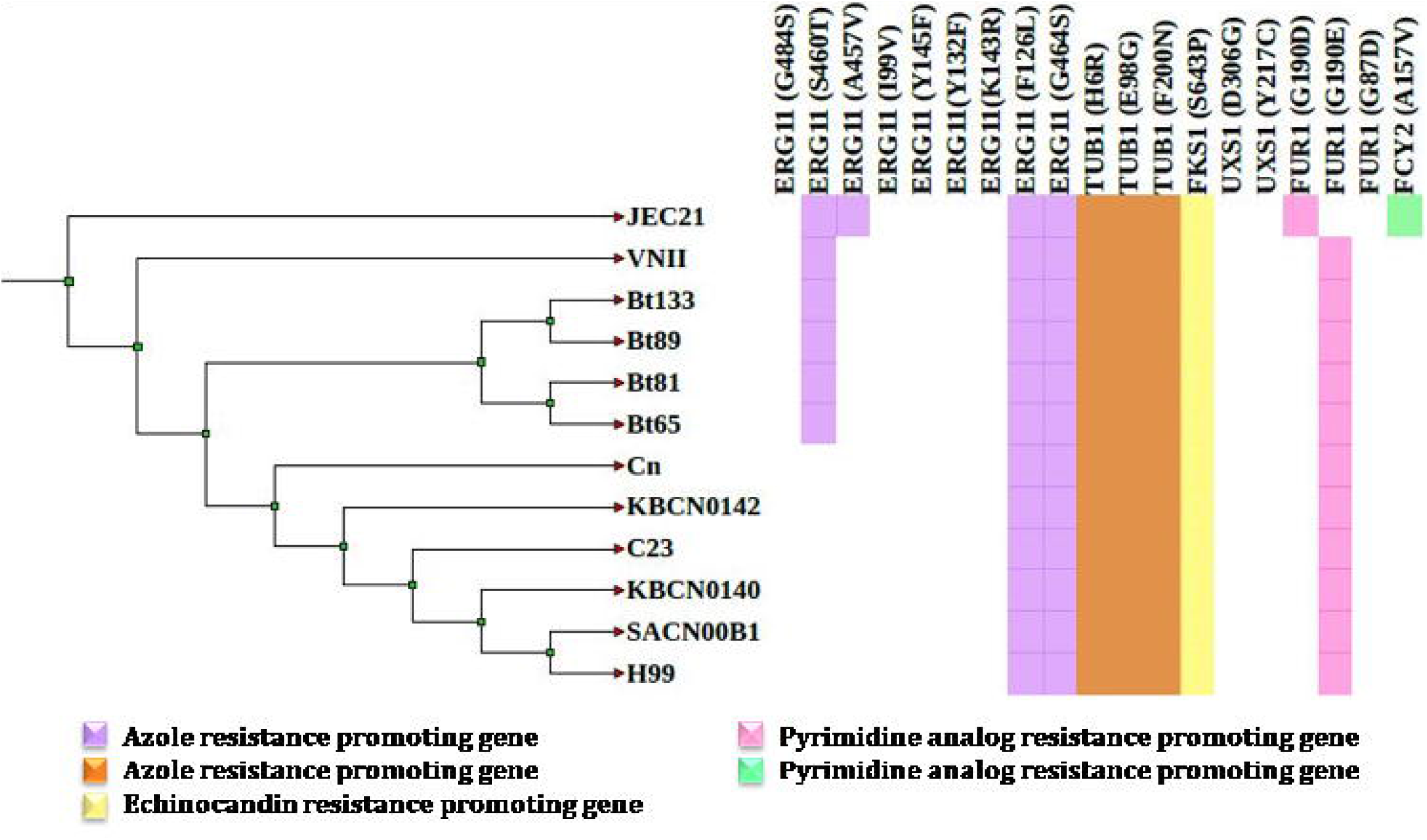
The preliminary orthologous analysis that shows the number of gene clusters (a), proteins (b) and singletons (c) present in each study strain.

**Table 2:** The SNPs present in AFR genes of *C. neoformans* study strains.

| **Genes** | **Residue in protein** | **Mutation detected differently than proposed in journals** |
| --- | --- | --- |
| **ERG11(9)** |  |  |
| Y145F | active site residue | - |
| G484S | active site residue | - |
| S460T | not an active site residue | - |
| A457V | not an active site residue | - |
| I99V | not an active site residue | - |
| Y132F | active site residue | - |
| K143R | not an active site residue | - |
| F126L | not an active site residue | F126S |
| G464S | not an active site residue | G464V |
| **FKS1 (1)** |  |  |
| S643P | not an active site residue | - |
| **TUB1 (3)** |  |  |
| H6Q | not an active site residue | H6R |
| E98 | not an active site residue | E98G |
| F200 | not an active site residue | F200N |
| **UXS1 (2)** |  |  |
| D306G | not an active site residue | - |
| Y217C | not an active site residue | - |
| **FUR1 (3)** |  |  |
| G190D | not an active site residue | - |
| G87D | not an active site residue | - |
| G190E | not an active site residue | - |
| **FCY2 (1)** |  |  |
| A157V | not an active site residue | - |

### 3.7. Pan-genome analysis and phylogenetic analysis using orthologous genes

The pan-genome analysis aids in understanding the genome architecture of a species. The preliminary orthologous gene analysis showed the number of gene clusters, proteins and singleton genes in each genome (figure 10a, 10b and 10c). The overall read statistics of the orthologous genome analysis are provided in Table 3. In figure 11, the gene clusters and their associated proteins have been designated as core, shell, and cloud. The orthologous analysis generates two data sets, one is the upset chart where the unique orthologous cluster and proteins present in the 12 genomes are independent of one another and the second is the occurrence chart where the unique orthologous cluster and proteins are dependent on each other. The occurrence chart and upset chart visualize the presence and absence of orthologous gene clusters and protein count across the compared study genomes. The unique clusters and proteins of the study genomes are derived from upset chart were the gene clusters and proteins are not correlated and demonstrated in table 4. The unique clusters and proteins of the study genomes are derived from occurrence chart were the gene clusters and proteins are correlated were demonstrated in table 5. The occurrence chart and upset chart provided a correlation between the gene cluster and protein count. From the occurrence chart of the gene cluster, it was depicted that there were about n=4676 clusters (73.6%) containing n=57595 proteins (77.3%) present in all the study genomes (figure 12, 13, 14). The pan-genome analysis also states the 4304 single-copy clusters (single-copy orthologous clusters) across the 12 genomes. The single copy clusters were categorized into biological process-associated (96.68%), molecular function-associated (2.23%) and cellular component-associated (1.08%) gene clusters (Figure 15; Table 6). The phylogenetic tree was constructed by the maximum likelihood method using the orthologous genes (figure 16). The midpoint was divided into two clades where the strain JEC21 (*C. neoformans var. neoformans*) was in **Clade1** and the rest in **Clade 2** occupied by the subspecies *grubii.* Clade 2 was further divided into 2 subclades where **subclade 1** has the environmental strain VNII (environmental strain) and **subclade 2** has clinical strains that was further categorized into 6 clusters. The gene expansion (gene duplication) and contraction (gene deletion) studies using the CAFE5 showed the addition of gene families (expansion) and contraction of gene families. The highest contraction of gene families was reported in the environmental strain VNII and the strains isolated from Australia showed mere stable expansion and contraction that were reported in figure 16. A pairwise heat map was generated based on the similarity of orthologous genes which shows that the strain Cn (clinical strain; sub sp. *grubii*) and VNII (environmental strain) have high similarity with JEC21 (clinical strain; sub sp. *neoformans*). The strain Bt65 (clinical strain) is similar to Bt81 (clinical strain) and Bt89 (clinical strain) is similar to Bt133 (clinical strain) (figure 17).

**Figure 11:**
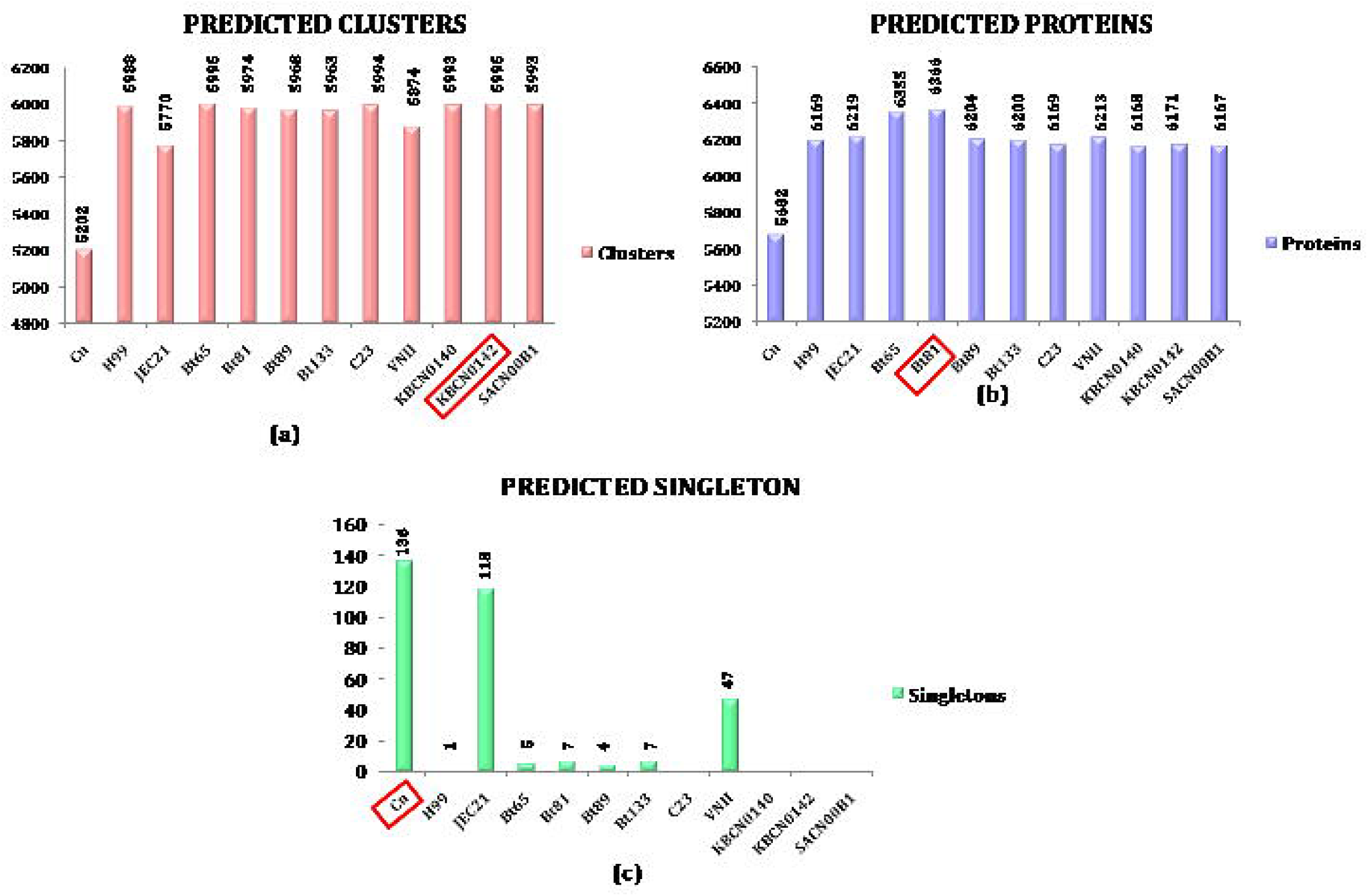
The pictorial representation of the overall orthologous clusters and proteins.

**Figure 12:**
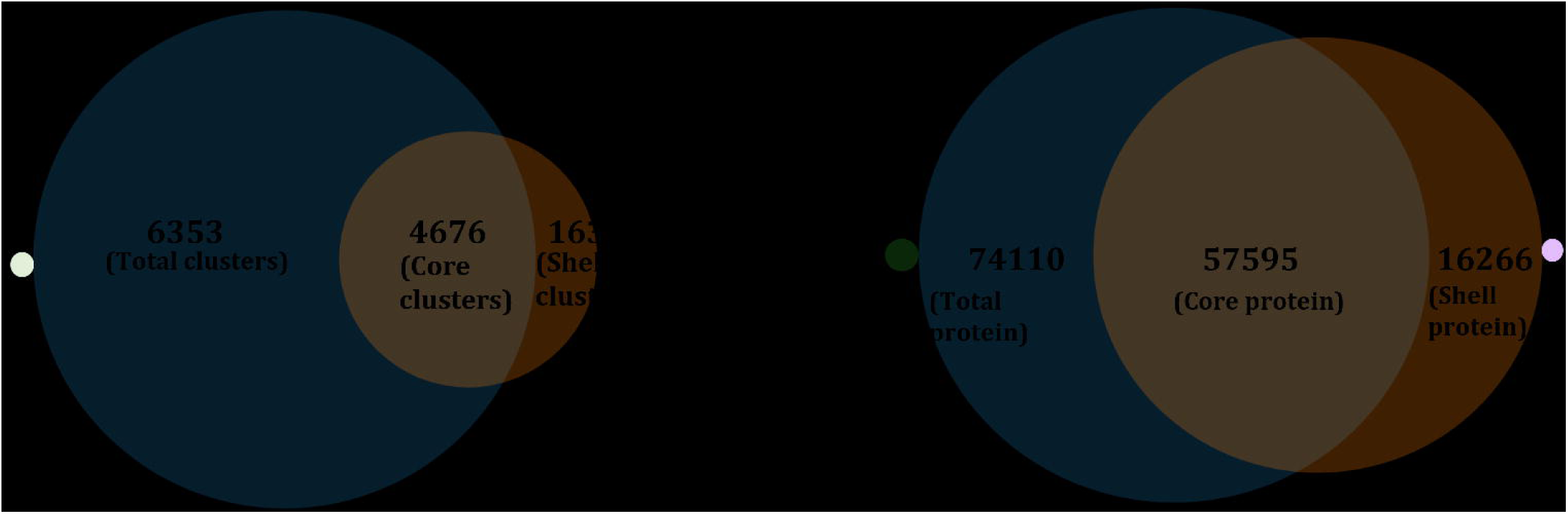
The pictorial representation of the occurrence chart that depicts the presence of orthologous gene clusters present and shared among the 12 genomes.

**Figure 13:**
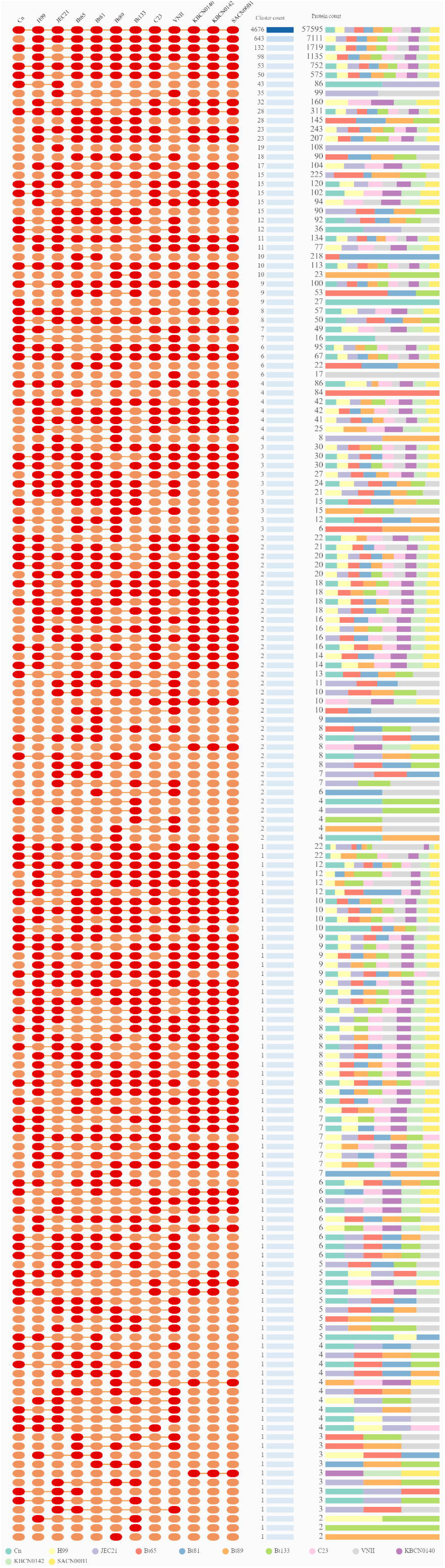
The pictorial representation of the occurrence chart that depicts the presence of overlapping gene clusters present and shared among the 12 genomes.

**Figure 14:**
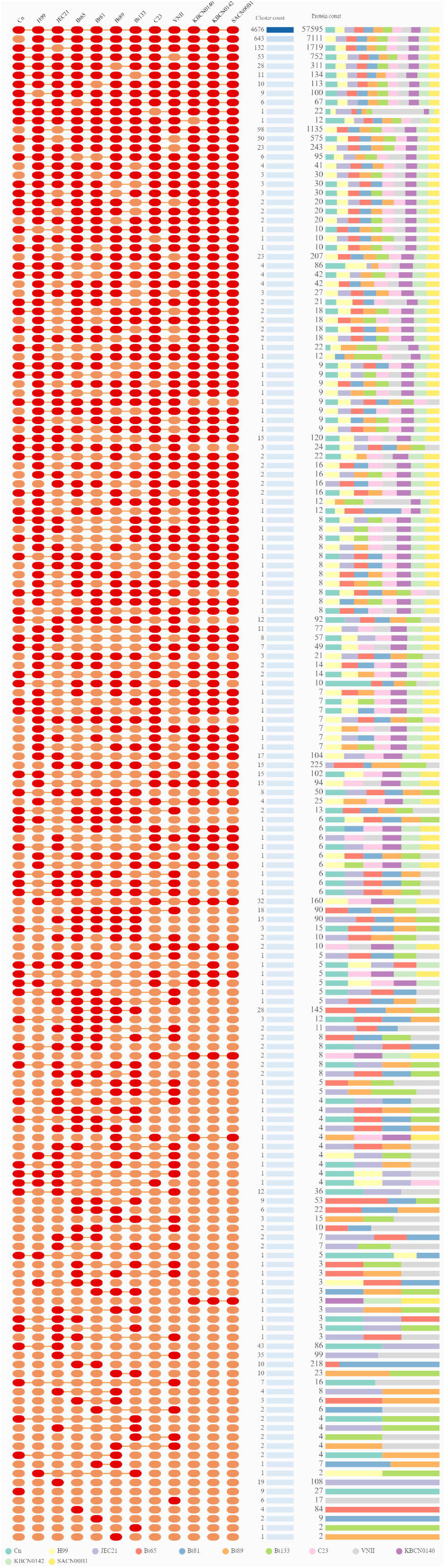
The pictorial representation of the occurrence chart that depicts the presence of proteins present and shared among the 12 genomes.

**Figure 15:**
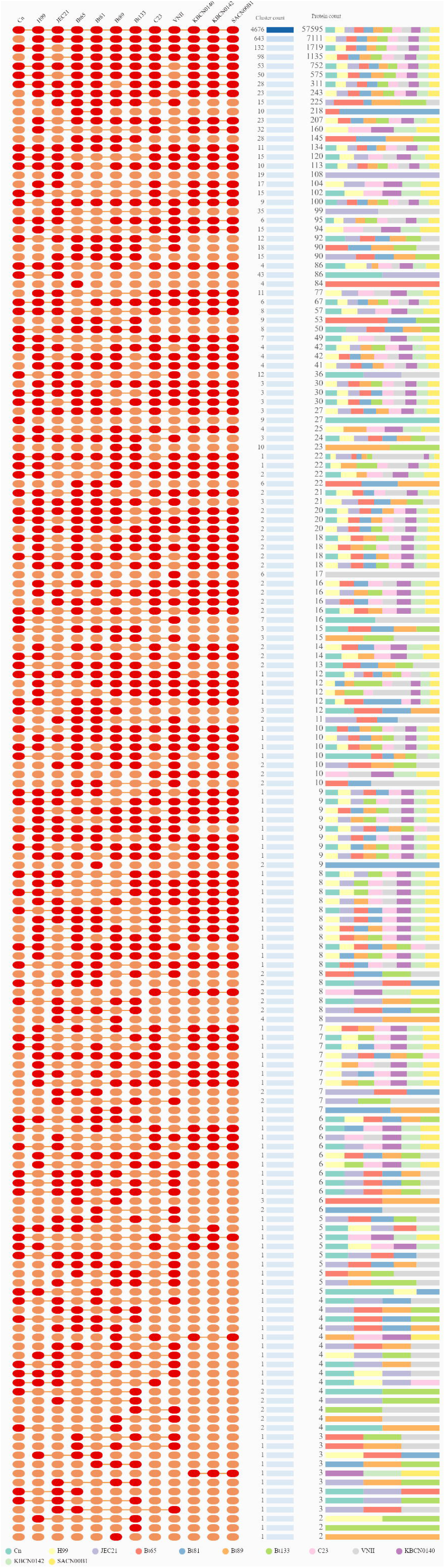
Single-copy gene clusters present in 12 genomes classified as cellular component genes (a), biological process gene (b) and molecular function genes (c)

**Figure 16:**
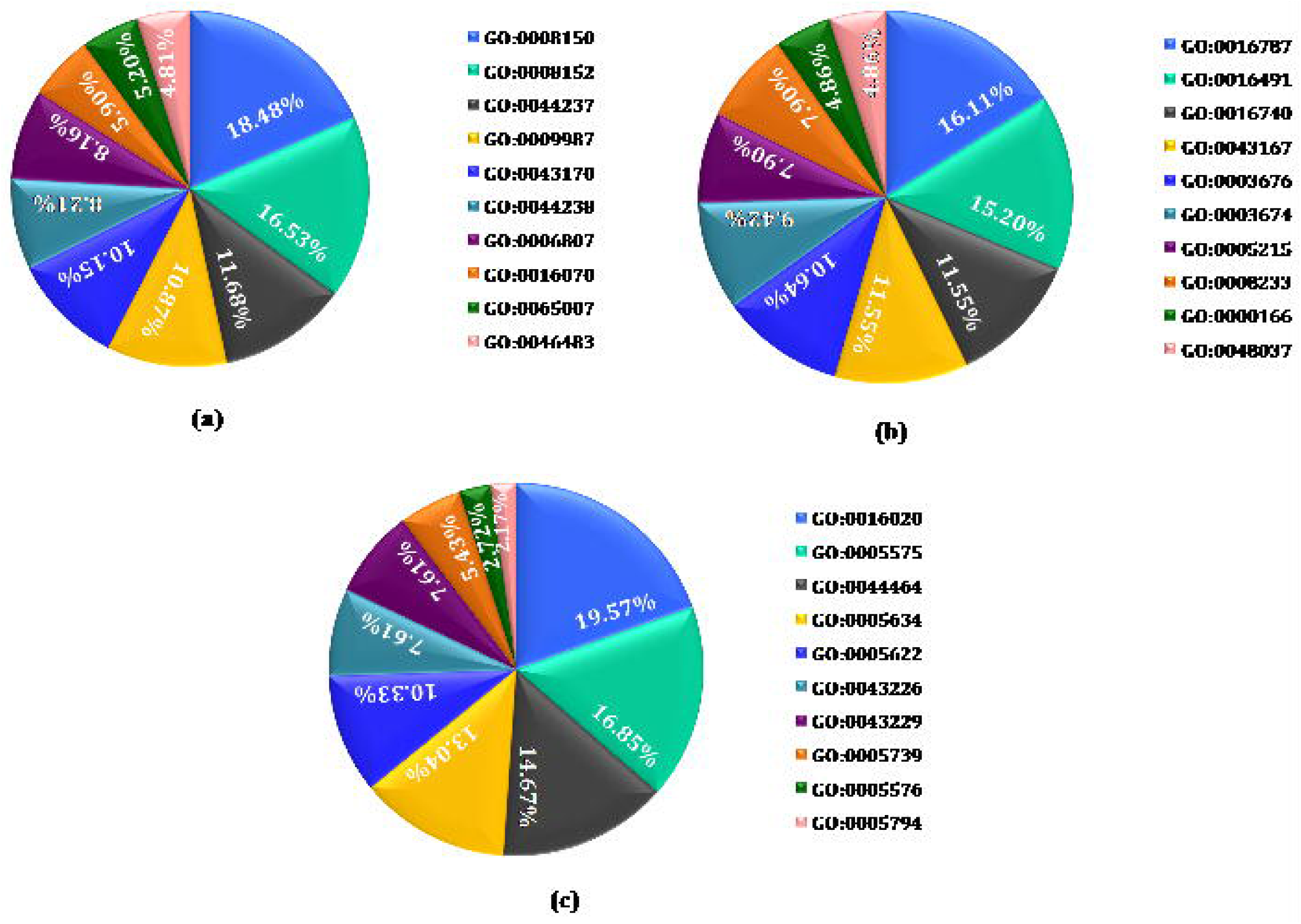
Phylogenetic tree based on orthologous genes: Orthologous gene-based phylogeny was constructed by the maximum likelihood method with CAFE5. The midpoint of the phylogenetic tree was divided into two major clades and further delineated into many clusters of *C. neoformans*.

**Figure 17:**
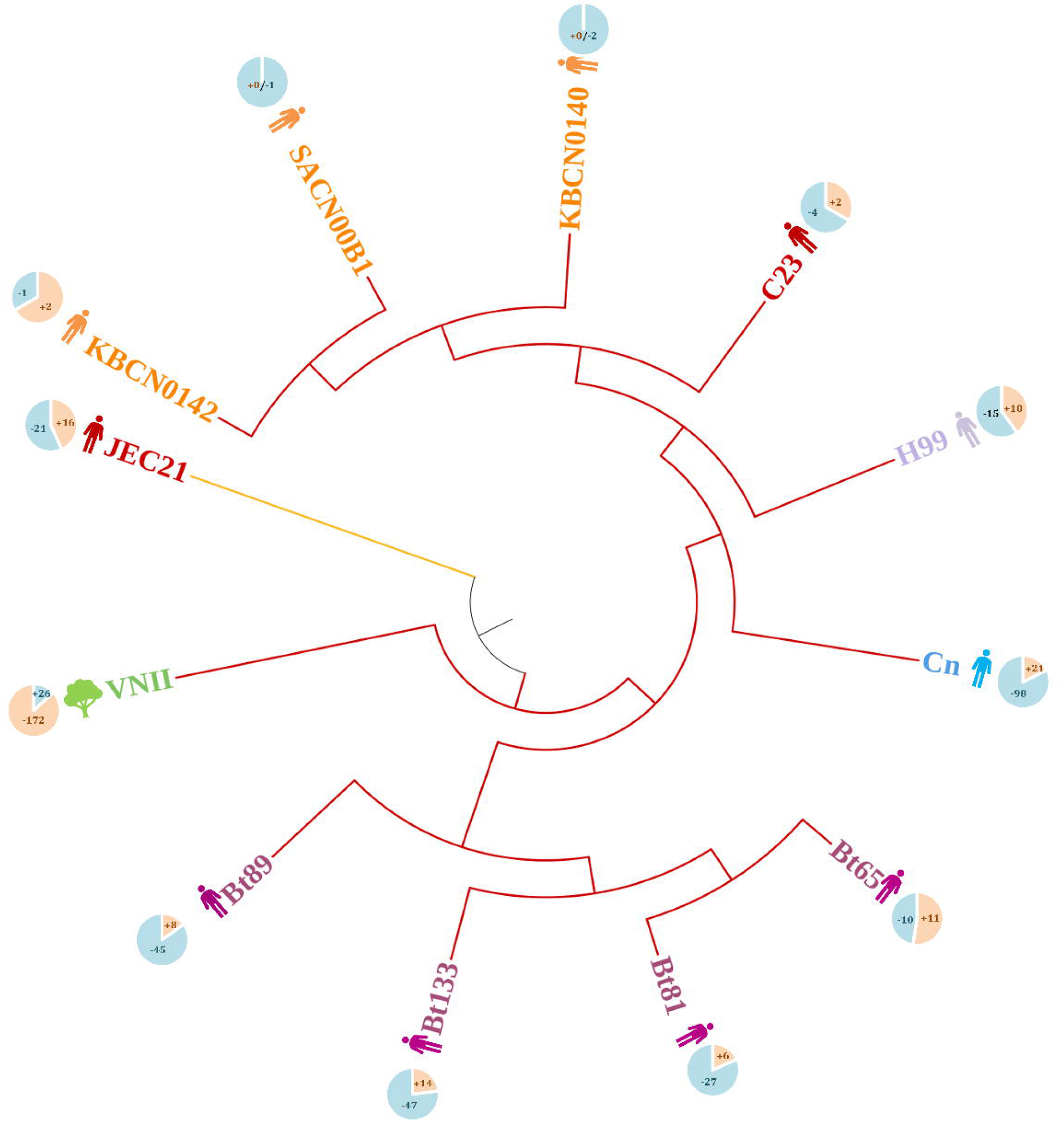
Pairwise heatmap: The intensity of the colour gradient shows the similarity between the species.

**Table 3:**
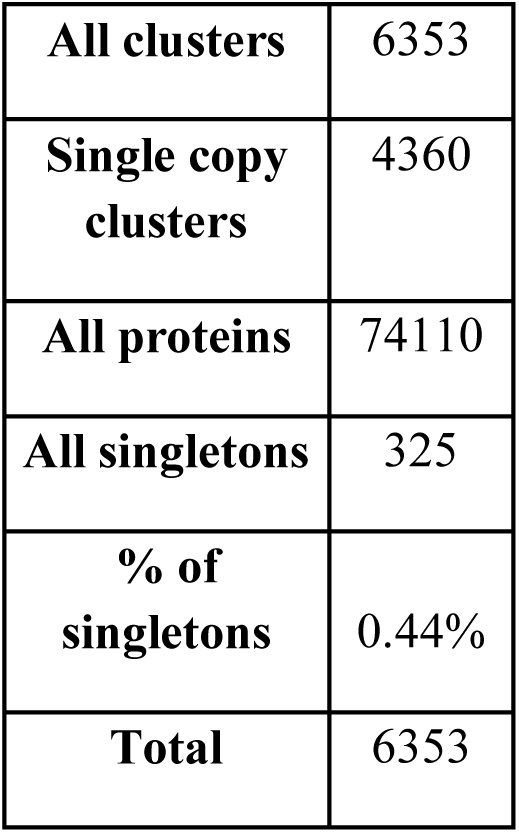
The overall read statistics of orthologous gene.

**Table 4:** The unique orthologous cluster and proteins present in the 12 genomes-Upset table where the unique clusters and protein count are independent.

| Strain | Unique cluster | Unique protein |
| --- | --- | --- |
| Cn | 9 | 9 |
| H99 | 89 | 89 |
| JEC21 | 19 | 19 |
| Bt65 | 4 | 4 |
| Bt81 | 2 | 2 |
| Bt89 | 1 | 1 |
| Bt133 | 1 | 1 |
| C23 | 88 | 88 |
| VNII | 6 | 6 |
| KBCN0140 | 87 | 87 |
| KBCN0142 | 89 | 89 |
| SACN00B1 | 87 | 87 |
| **Total** | **482** | **482** |

**Table 5:** The unique orthologous cluster and proteins present in the 12 genomes-Occurrence table where the unique clusters are correlated with the protein count.

| **S.no.** | **Strain name** | **Unique clusters** | **Unique proteins** |
| --- | --- | --- | --- |
| 1. | JEC21 | 19 | 108 |
| 2. | Cn | 9 | 27 |
| 3. | Bt65 | 4 | 84 |
| 4. | Bt81 | 2 | 9 |
| 5. | VNII | 6 | 17 |
| 6. | Bt133 | 1 | 2 |
| 7. | Bt89 | 1 | 2 |
| **Total** |  | **42** | **249** |

**Table 6:**
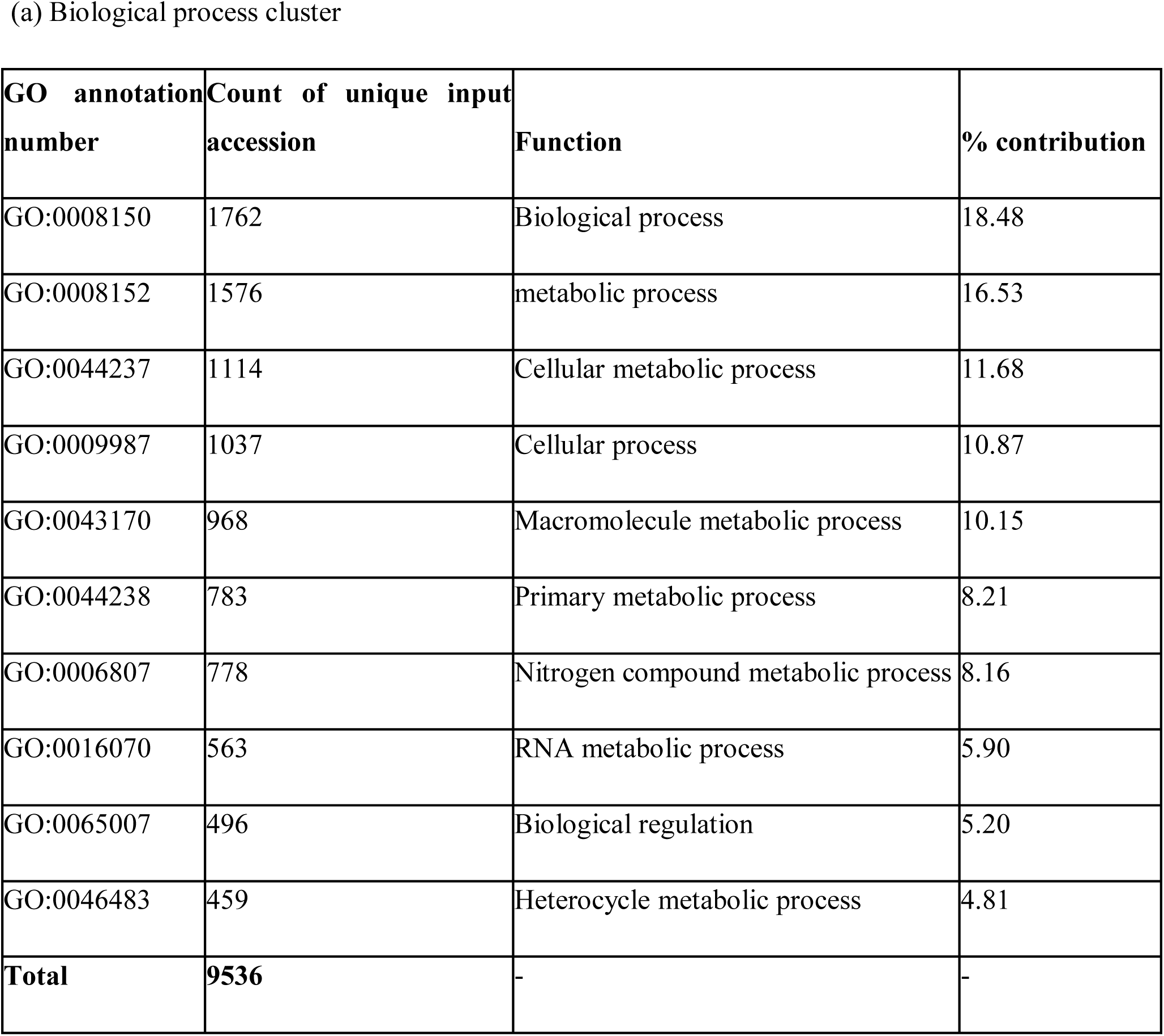
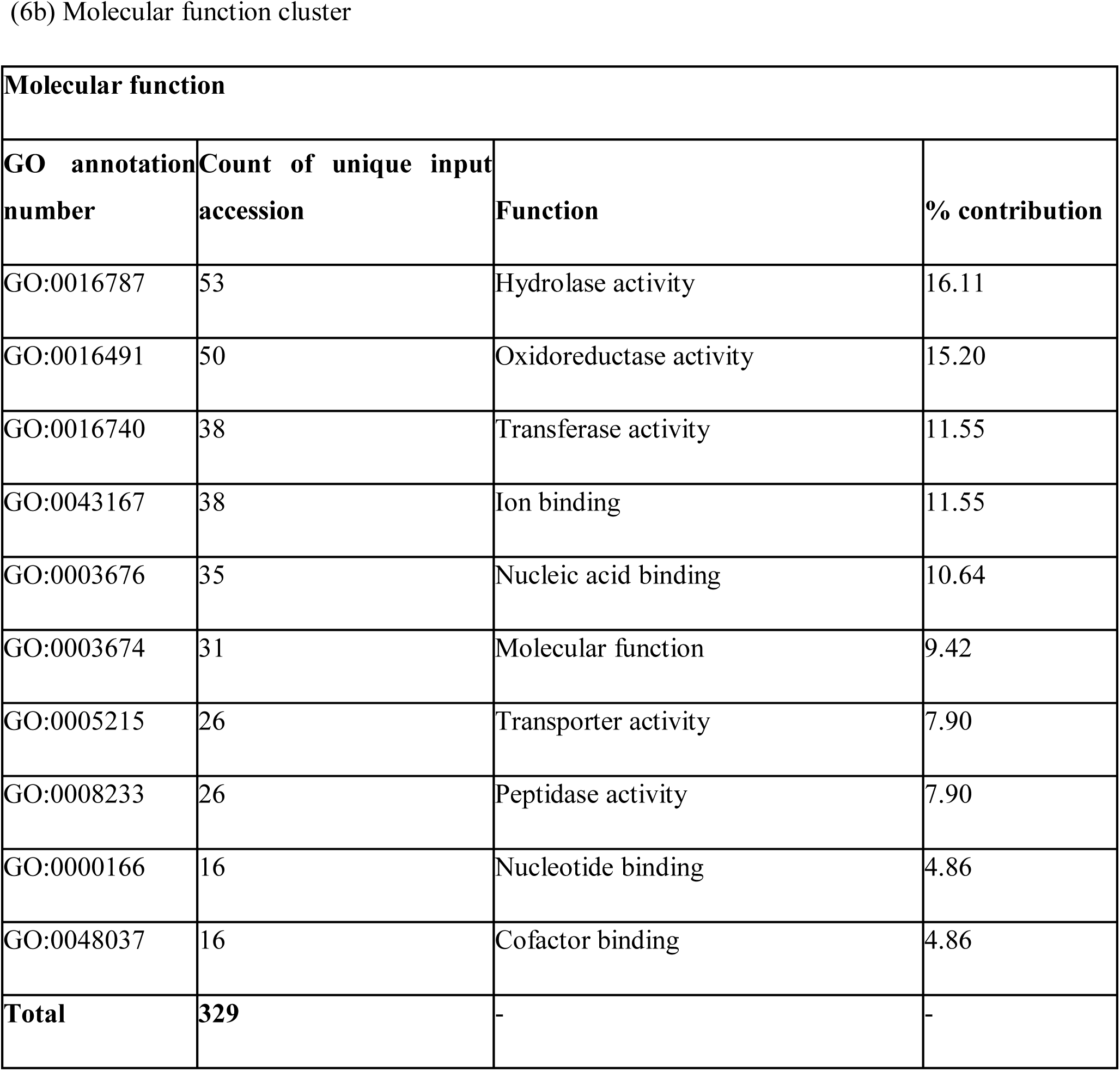
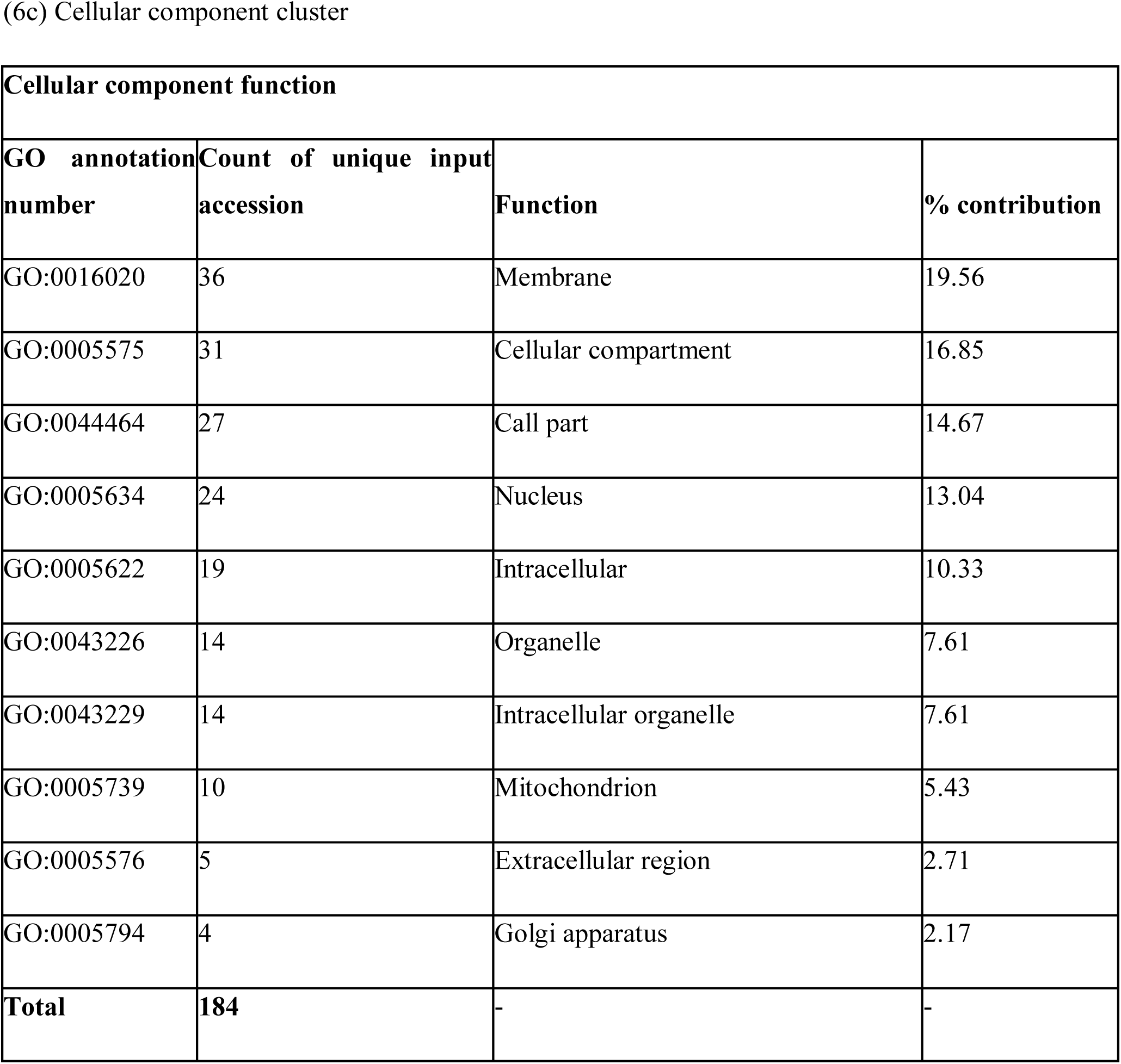
The single copy number genes present across the 12 genomes.

(a) Biological process cluster
| **GO annotation number** | **Count of unique input accession** | **Function** | **% contribution** |
| --- | --- | --- | --- |
| GO:0008150 | 1762 | Biological process | 18.48 |
| GO:0008152 | 1576 | metabolic process | 16.53 |
| GO:0044237 | 1114 | Cellular metabolic process | 11.68 |
| GO:0009987 | 1037 | Cellular process | 10.87 |
| GO:0043170 | 968 | Macromolecule metabolic process | 10.15 |
| GO:0044238 | 783 | Primary metabolic process | 8.21 |
| GO:0006807 | 778 | Nitrogen compound metabolic process | 8.16 |
| GO:0016070 | 563 | RNA metabolic process | 5.90 |
| GO:0065007 | 496 | Biological regulation | 5.20 |
| GO:0046483 | 459 | Heterocycle metabolic process | 4.81 |
| **Total** | **9536** | - | - |

(6b) Molecular function cluster
| **Molecular function** |  |  |  |
| --- | --- | --- | --- |
| **GO annotation number** | **Count of unique input accession** | **Function** | **% contribution** |
| GO:0016787 | 53 | Hydrolase activity | 16.11 |
| GO:0016491 | 50 | Oxidoreductase activity | 15.20 |
| GO:0016740 | 38 | Transferase activity | 11.55 |
| GO:0043167 | 38 | Ion binding | 11.55 |
| GO:0003676 | 35 | Nucleic acid binding | 10.64 |
| GO:0003674 | 31 | Molecular function | 9.42 |
| GO:0005215 | 26 | Transporter activity | 7.90 |
| GO:0008233 | 26 | Peptidase activity | 7.90 |
| GO:0000166 | 16 | Nucleotide binding | 4.86 |
| GO:0048037 | 16 | Cofactor binding | 4.86 |
| **Total** | **329** | - | - |

(6c) Cellular component cluster
| **Cellular component function** |  |  |  |
| --- | --- | --- | --- |
| **GO annotation number** | **Count of unique input accession** | **Function** | **% contribution** |
| GO:0016020 | 36 | Membrane | 19.56 |
| GO:0005575 | 31 | Cellular compartment | 16.85 |
| GO:0044464 | 27 | Cell part | 14.67 |
| GO:0005634 | 24 | Nucleus | 13.04 |
| GO:0005622 | 19 | Intracellular | 10.33 |
| GO:0043226 | 14 | Organelle | 7.61 |
| GO:0043229 | 14 | Intracellular organelle | 7.61 |
| GO:0005739 | 10 | Mitochondrion | 5.43 |
| GO:0005576 | 5 | Extracellular region | 2.71 |
| GO:0005794 | 4 | Golgi apparatus | 2.17 |
| **Total** | **184** | - | - |

## 4. Discussion

In this study, we sequenced and assembled a genome of *C. neoformans* (Cn) collected from the Microbiology lab of Trichy Medical College, Tamil Nadu, India sourced from cryptococcal meningitis patient. Together with the already available 11 genomes of the species, we compared all 12 genomes and deciphered the phylogenics, resistome, virulome and pangenome of these cryptococcal strains. The genome size of all the study strains is 18.9 to 19.1 Mb and the GC contents ranged between 48 to 48.5%.

The ISHAM ITS barcoding helped to confirm the subspecies of the study strain Cn to be *grubii* (100% identity) which is in correlation with the constructed phylogenetic tree. The Cn study strain’s subspecies identification also confirms that it is a member of the serotype A. Due to the lack of 100% matching sequences in the ISHAM database, we anticipate that the strain Cn might belong to a new ST type.

The phylogenetic tree constructed using the MLST genes and the orthologous genes shows similar results, especially on the strains from Botswana, South Africa. Except for the strain JEC21(California, USA), all the other strains belong to the same subspecies *grubii* the strains of USA, Australia and India showed diversities in the clades. The strains from Botswana, South Africa are present in the same subclade showing the convergence in their genome. Sub-Saharan Africa has been the hub for Cryptococcal infection and the Botswana belongs to the southern part of Sub-Saharan Africa [12]. The genetic convergence is obvious in this part due to several factors such as strong selective pressures, high transmission rate, limited gene flow and host adaptation.

Our manual method of retrieving virulence genes from literature and BLAST analysis revealed the presence of 99.28% of virulence genes in H99. In various reported studies virulence nature of H99 was well demonstrated using animal models [45,46]. In addition, our recent demonstration of the disseminated cryptococcosis mice model also revealed the high virulent potency of H99. Confirming our manual method as an ideal one for virulome analysis, however, the creation of a virulome database is the need of the hour. The virulome analysis depicts the presence of stress tolerance genes, which are pleiotropic and contribute to virulence regulation. The ability of *Cryptococcus* sp., to adapt to the host environment depends on how well it can maintain metabolic homeostasis under stress. The upregulation of carbohydrate transport, lipid transport, and autophagy, indicates that there is a significant need for the uptake of nutrients and energy production after host infection and phagocytosis. The *Cryptococcus* sp. has a set of autophagy genes that are essential for sustaining cellular integrity, homeostasis and survival (stress tolerance). Some of the metabolic pathways such as the pentose phosphate pathway which has been regulated by ATP were also upregulated under nitrosative stress conditions. This serves as an example that the metabolic pathways were likely to be induced by stress conditions. A recent study demonstrated that the transcription factor CIR1 (master iron regulator) and RIM101 integrate iron homeostasis with a multifarious of other functions including pH sensing, nutrient and stress signaling pathway, virulence factor elaboration and cell wall biogenesis [15–17].

The study strains C23, Bt89, and Bt133 was found to lack the major virulence factor URE1, which serves to breach the BBB and host immune system. Except Cn, H99 and JEC21 all the other strains lack the gene *LAC1* that contributes majorly to the melanin production, biofilm production and inhibition of phagocytic uptake. This is also concurrent with our previous study, in which the Cn was found to have a dominant capsule, melanin production and biofilm formation [47].

While comparing the virulome of the study strain Cn and the reference genome (H99), *ATG12*, *ATG16, CHS4*, Alkaline phosphatase, *UGD1*, *APP1*, *BOS1* and *RAS2* were found to be absent in the study strain which contributes to immune evasion. While comparing the virulome of the reference strain and the environmental strain, *ATG4*, *ATG9*, *ATG12*, *ATG14*, *ATG18*, *ATG26*, *CBP1*, *XPT1*, *CHS4*, *CHS6*, *CHS8*, *CDA3*, *URE1*, phosphatidic acid phosphatase, *SIP4*, *CAN1*, *FRR1*, *SHO1*, *CLN1*, *SEC61B*, *SEC16*, *LAC1*, copper transport protein 86, potassium transport protein, *PDK1*, Outer cell wall protein RRT8, Yeast cell wall synthesis Kre9/Knh1-like N-terminal domain-containing protein, TatD DNase family Scn1, *CAN2*, homoserine kinase, Homoserine O-acetyltransferase, α-amylase, Alkaline phosphatase, 2-Methylcitrate synthase, *UGD1*, RNaseIII, Inositolphosphorylceramide synthase subunit Kei1-domain-containing protein, *BIM1*, *VMA21*, *UGT1*, *BOS1*, *RUB1*, *SCH9*, *RAS2* were found to be absent in the environmental strain that has protease, urease, laccase, autophagy-related and cellular function.

All the 12 isolates showed most of the major VRGs (above 80%) to be found in their genome. The environmental factors such as ecological niches, temperature and host immune responses enhance the potential of Cryptococcal virulence. The soil and bird droppings are the nutrient-rich environment supporting growth and enhancing its virulence potential. The enhancement of virulence in nutrient-rich environments is due to the conserved cAMP/ PKA signaling pathway that plays a crucial role in nutrient sensing and aids in the regulation of virulence genes [48]. The ability of *Cryptococcus* spp. to thrive at 37 °C (normal human body temperature) is crucial for its pathogenicity. The interaction of *Cryptococcus* spp. with the host immune system is critical for its virulence as it can modulate the immune response pathways, particularly inhibiting the inflammasome pathway, which is essential for effective antifungal immunity[49]. This immunomodulatory activity helps the *Cryptococcus* spp. to survive and multiply within the macrophages, further enhancing its virulence. The *Cryptococcus* spp. ability to adapt to various ecological niches, coupled with its capacity to modulate the host immune responses, underscores the importance of environmental influences on its pathogenicity.

Even though all the strains contained the AFR genes with certain mutations, most of the mutational sites do not belong to the active site binding pocket. As additional supporting data, the *in vitro* investigation on the study strain and the reference strain (H99) demonstrated that both the strains were susceptible to amphotericin B (polyenes) and fluconazole (azole). With this knowledge, it was anticipated that a mutation that occurs outside the active site binding pocket might not be the cause of AFR. This comparative genomic study of *C. neoformans* on virulence and AFR gives insights into the potent targets for the development of novel antifungal drugs.

Approximately 70 % of Cryptococcal genomes have conserved functional elements (genes involved in signaling and metabolic pathways) that are pivotal for survival and enhanced pathogenicity. The investigation of these conserved functional elements helps to develop more effective therapeutic strategies to combat this opportunistic pathogen.

The pan-genome analysis of the *Cryptococcus neoformans* gives various insights into the orthologous gene and protein levels. The number of gene clusters and proteins are not necessarily correlated. The strain KBCN0142 has the highest number of clusters (n=5995) and the strain Bt81 has the highest number of proteins (n=6366). This might be due to the fact of high redundancy, high similarity among the proteins, stringent clustering parameters, and the presence of paralogous genes. The presence of singleton proteins in the genomes shows that they are essential proteins that appear unique within the dataset and don’t have any closely related homologs. The strain Cn has the highest number of singletons up to 136. The singleton showcases the unique protein function and evolutionary significance. The orthologous analysis performed demonstrated the intersection of gene clusters and proteins; unique clusters and unique proteins of each genome. Though there were unique clusters and unique proteins present in all the 12 study strain genomes, as represented by the upset charts of cluster count and protein, the occurrence chart where the gene clusters are correlated with the protein shows the absence of a unique cluster or protein in H99, C23, KBCN0140, KBCN0142 and SACN00B1. The single-copy clusters depict the group of orthologous genes that contains exactly one copy of a gene in the cluster of the study genome. Such clusters are exceptionally important in comparative genomics and evolutionary study as they represent the conserved genes that reduce the noise from gene duplication or loss events in the compared species. The single-copy gene clusters were categorized based on functional nature as a biological process, molecular process and cellular component. Among them, the biological process-related GO cluster contributes to a maximum of 96.69% (n=17262), molecular function contributes to about 2.23% (n=398) and finally, the cellular component contributes to about 1.08% (n=193). This shows that the biological process-related genes are more conserved across the species. Though the strain JEC21 belongs to the subspecies *neoformans* and the strain Cn belongs to sub sp. *grubii* the pairwise heat map shows high similarity between them which might be because both the subspecies were descendants of the same ancestor. The phylogenetic tree constructed with the orthologous genes was considered to show more evolutionary relatedness than the tree constructed with the housekeeping genes. The reason behind the statement is the comprehensive representation of the genome, robustness to homoplasy, reduced noise of gene duplication or gene loss and improved alignment quality.

These factors reflect the true evolutionary relationships among the species. In this study, the phylogenetic tree constructed using the MLST genes and orthologous genes are merely similar. The MLST genes show that the South African strains are categorized in a single clade whereas the Pan-genome phylogenetic tree shows that the strains from South Africa and Australia are also in convergence respective to their isolation regions. The gene expansion and contraction studies showed that there was expansion (additional gene copies have been gained in their lineage) or contraction (number of gene copies lost in the lineage) among the genomes of the strains being isolated from the same region. The strains from Australia alone showed mere stable gene expansion and contraction due to the presence of stable gene families (the gene families in the species are stable concerning other species showing no significant changes in size due to duplication or loss events) or equal evolutionary changes (either the expansion/contraction occurring uniformly/ not at all occurring across all the species being analyzed leading to unique changes in a single lineage).

## 5. Conclusion

In conclusion, this was the first study to report the complete genome sequence of a *Cryptococcus* sp. from India and to study comparative genomics of strains from different geographical locations worldwide. Our study revealed that phylogenetic tree construction using orthologous genes yields more accurate evolutionary relationships compared to MLST. Furthermore, virulome analysis identified a high presence of major VRGs in Cn and H99 strains, consistent with *invivo* virulence assays. Resistome analysis detected major AFR genes, but mutations were absent in the active site binding pocket, aligning with *in vitro* MIC tests. The study suggests, the need for a comprehensive study which includes, virulome analysis, active site prediction, and *invitro* tests to confirm the drug resistance in fungal species, very specifically to the strains that lack the critical antibiotics breakpoints. Pan-genome analysis showed gene expansion and contraction across strains. Genome plasticity (gene expansion and contraction) is a critical factor of *Cryptococcus neoformans* sub-sp. in its adaptability, virulence and ability to resist antifungals. Deciphering these mechanisms provides insights into this pathogen evolution and suggests potential targets for therapeutic interventions. In summary, the findings on virulome and resistome profiles shed light on the pathogenicity of different strains which is crucial for understanding disease management. The study advocates the need for bioinformatic tools for resistome and virulome analysis.

## Supporting information

Supplementary figure 1: Agarose gel electrophoresis of isolated cryptococcal DNA: A-represents the HIND III digested lambda ladder and B-represents

Supplementary figure 2: Tape station profiling: Fragment size distribution analysis of the prepared library

## Acknowledgment

We thank SASTRA Deemed University for providing the research facilities and infrastructure.

## List of supplementary figures

**Supplementary figure 1:**
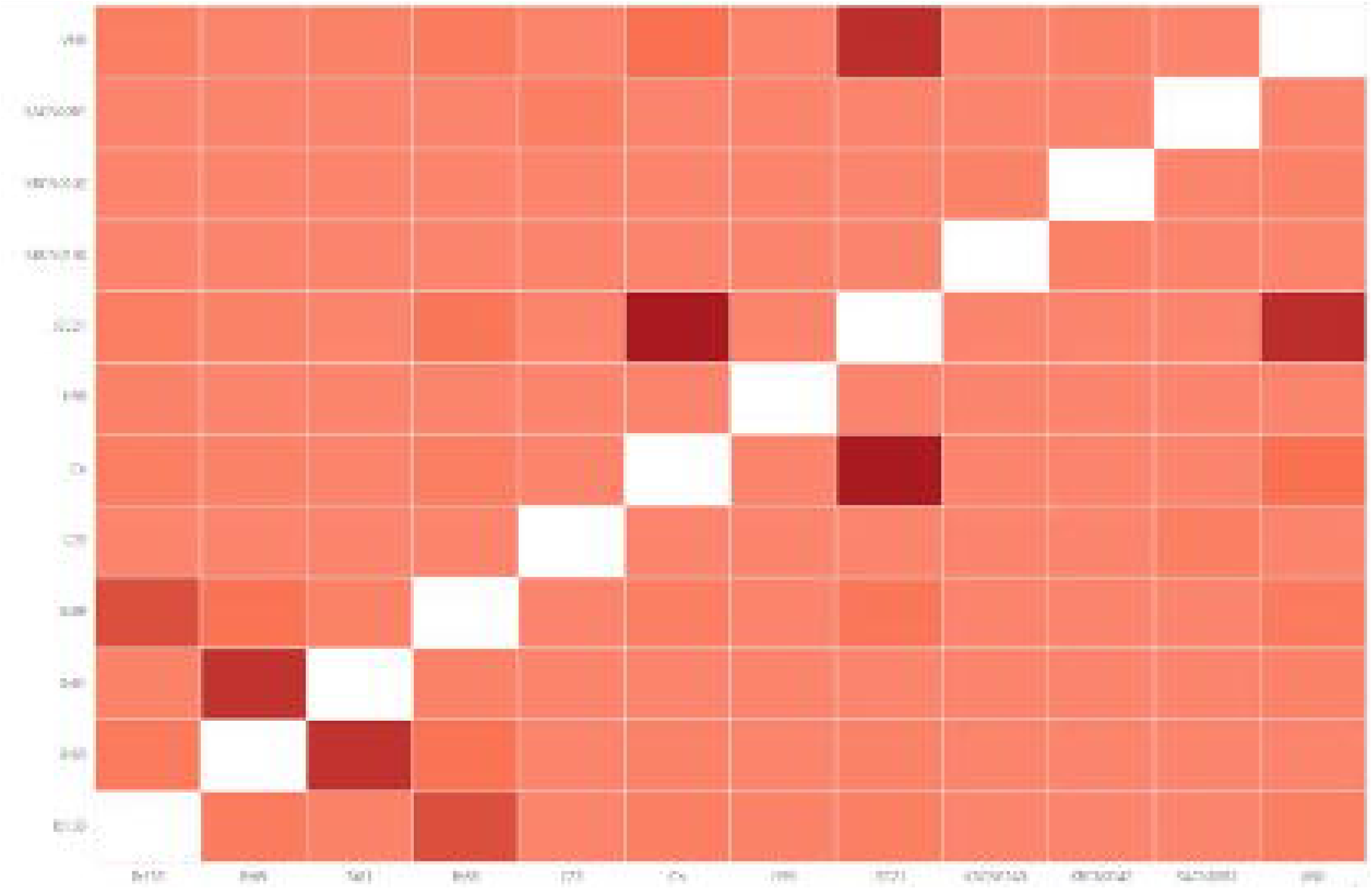
Agarose gel electrophoresis of isolated cryptococcal DNA: A-represents the HIND III digested lambda ladder and B-represents the isolated DNA of Cn strain.

**Supplementary figure 2:** Tape station profiling: Fragment size distribution analysis of the prepared library.

