## Supplementary figures and images for "Hybrid Genome Sequence of *Cryptococcus neoformans* of Indian origin and Comparative Genome Analysis"

### Supplementary figure 1: Agarose gel electrophoresis of isolated cryptococcal DNA: A-represents the HIND III digested lambda ladder and B-represents

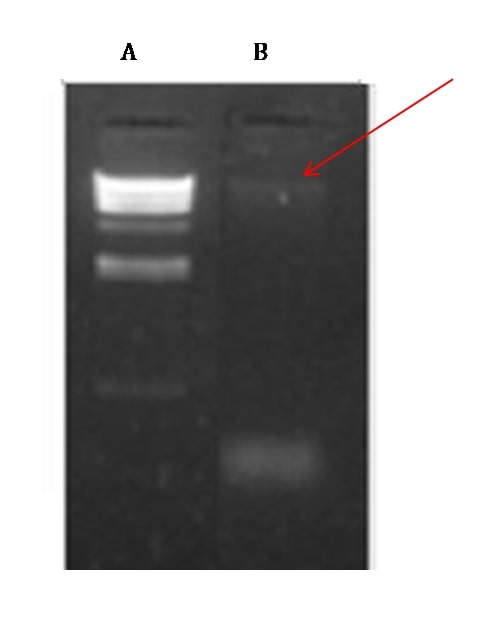

### Supplementary figure 2: Tape station profiling: Fragment size distribution analysis of the prepared library

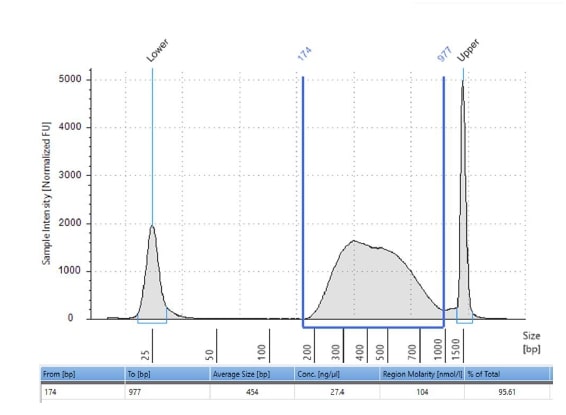
